# Disruption of the hypothalamic orexin system links SARS-CoV-2 infection to persistent cortical neuronal pathology

**DOI:** 10.64898/2026.01.22.701182

**Authors:** Gun Young Yoon, Young-Chul Jeong, Ji-Hyun Choi, Yoon Ha, Se Yeon Seo, Keun Bon Ku, Do Yeon Kim, Woo Yeon Hwang, Gi Uk Jeong, Dae-Gyun Ahn, Kyun-Do Kim, Je-Keun Rhee, Won-Ho Shin, Young-Chan Kwon

## Abstract

Long COVID frequently presents with persistent neurological symptoms, including cognitive impairment, fatigue, and sleep disturbances; however, its underlying mechanisms remain unclear. Here, we show that SARS-CoV-2 infection induces lasting cortical neuronal injury and hypothalamic orexin (hypocretin) dysfunction *in vivo*. In K18-hACE2 and wild-type BALB/c mice, viral RNA persisted in the brain and coincided with focal loss of Neuronal Nuclei (NeuN)-positive cortical neurones beyond acute infection. SARS-CoV-2, but not the influenza A virus, triggered rapid and sustained suppression of hypothalamic orexin expression, defining a virus-specific neuropathological signature. Considering the downregulation of orexin and focal cortical NeuN expression, we showed that augmenting orexin signalling using recombinant orexin-A/B restored NeuN expression *in vitro* and *in vivo*. Overall, these findings identify the orexin system as a selective neural vulnerability to SARS-CoV-2 and define orexinergic circuit disruption as a mechanistic axis underlying the neurological manifestations of Long COVID.

## Main

Long COVID, defined as persistent symptoms lasting beyond four weeks after acute SARS-CoV-2 infection, affects approximately 10–30% of convalescent individuals^1,2^. Neurological and neuropsychiatric manifestations—, such as cognitive impairment, sleep disturbance, fatigue, and attention—deficits, represent some of the most debilitating outcomes documented across longitudinal cohorts^3–5^. Large-scale neuroimaging studies have revealed that SARS-CoV-2 infection induces long-lasting alterations in cerebral structure and function, including progressive cortical thinning, grey matter loss, and disruption of functional connectivity^6–8^. These changes correlate with persistent cognitive symptoms, indicating sustained disruption of neural circuits beyond the acute phase^5–7^.

Several mechanisms have been proposed to explain these effects. Neuropathological studies have revealed viral RNA and proteins in neurones, astrocytes, and microglia, confirming central nervous system (CNS) tropism^9–11^. Robust systemic immune activation is associated with cognitive symptoms and blood brain barrier breakdown^9,12^. However, the current neuroinflammatory models do not fully account for these key observations. Neurological sequelae often persist after overt inflammatory signatures wane, and all neurotropic viruses do not produce the same phenotype despite comparable immune activation^13,14^. This dissociation suggests that SARS-CoV-2 targets specific neural populations or homeostatic circuits whose dysfunction sustains neurological deficits.

Among these circuits, orexin (hypocretin), a neuropeptide that couples vigilance with energy balance, is produced by a small, anatomically restricted population of lateral hypothalamic neurones that nevertheless exerts outsized control over wakefulness and attention^15–17^. Selective loss of orexin neurones causes narcolepsy type 1, a disorder that shares key features with Long COVID, including sleep fragmentation and cognitive dysfunction^18,19^. Beyond regulating arousal, orexin promotes neuronal survival under oxidative stress^20^ and limits neuroinflammation^21^. However, whether orexin signalling directly regulates cortical neuronal integrity remains unclear. Given the established roles of the orexin system in homeostatic control and neuronal protection^15–17,21,22^, we hypothesised that SARS-CoV-2 infection compromises this critical network, potentially reducing circuit resilience^23,24^.

In this study, we identified two convergent *in vivo* signatures of SARS-CoV-2 neuropathology: focal, heterogeneous loss of cortical NeuN immunoreactivity persisting beyond the acute phase, and selective suppression of hypothalamic hypocretin/orexin (*Hcrt*) expression during early infection. Using comparative viral models, we showed that this specific combination was intrinsic to SARS-CoV-2 and was not recapitulated by the influenza A virus, despite showing comparable neuroinvasion. Furthermore, administering recombinant orexin-A/B partially restored cortical NeuN expression during infection. These findings identify the orexinergic system as a selective target of SARS-CoV-2 and suggest that disrupted orexin signalling may compromise cortical neuronal integrity during infection, thereby providing a new framework for understanding the persistent neurological consequences of COVID-19.

## Results

### Persistent cortical neuronal dysfunction and prolonged brain viral RNA in SARS-CoV-2 infected K18-hACE2 mice

K18-hACE2 mice are an established model for severe COVID-19, recapitulating the key features of CNS involvement, including viral neuroinvasion and neuroinflammation.^25,26^. We first confirmed the brain permissiveness following intranasal infection with a lethal dose (2 × 10^4^ PFU) of SARS-CoV-2, which led to robust NP detection in the olfactory bulb and cerebral cortex at 6 days post-infection (dpi) (Extended Data Fig. 1). To model long COVID, we employed a low-dose (50 PFU) infection paradigm, enabling long-term assessment for up to 90 dpi (Fig. 1a).

**Fig. 1.**
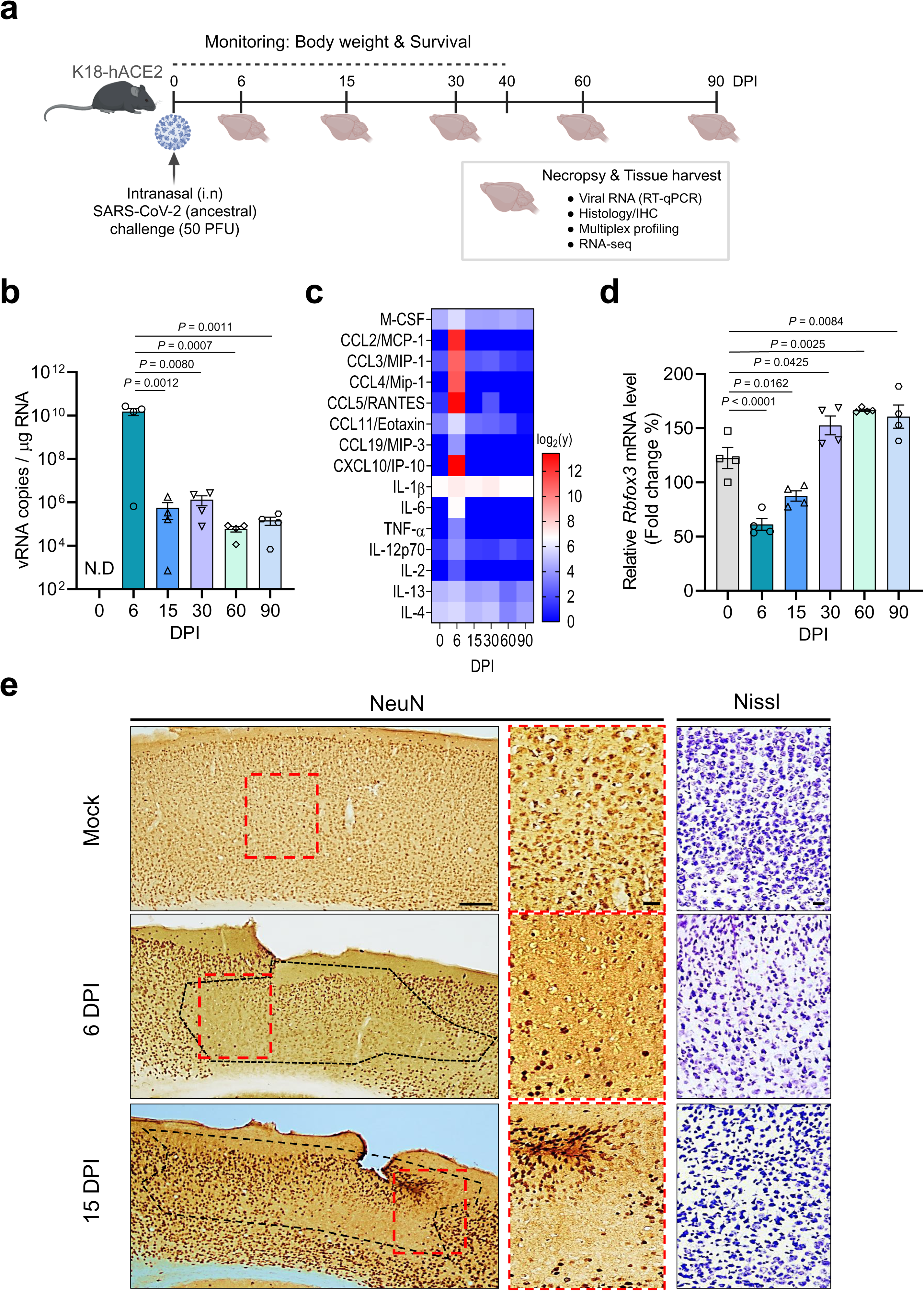
SARS-CoV-2 infection causes transient *Rbfox3* transcriptional suppression but persistent focal loss of cortical NeuN immunoreactivity in K18-hACE2 mice. **a**, Schematic of the experimental timeline. K18-hACE2 mice were intranasally challenged with 50 plaque forming units (PFU) of ancestral SARS-CoV-2 and monitored for up to 90 days post-infection (dpi). Tissues were collected at the indicated time points for viral load analysis, multiplex cytokine profiling, histopathology, and bulk RNA sequencing (n = 3-5 mice per group per time point). **b,** Time-course quantification of viral RNA burden in whole-brain homogenates using RT-qPCR, presented as viral RNA copies per µg of total RNA. **c,** Heat map of cytokine and chemokine protein levels in brain lysates at the indicated time points, measured using a multiplex array. The data represent log_2_-transformed concentrations (pg mg ¹ protein). Values <LOD were set to 1.0 for log transformation. **d,** Time-course quantification of *Rbfox3* (NeuN) mRNA expression in whole-brain lysates, presented as fold change relative to mock controls. **e,** Representative NeuN immunohistochemistry (left) and Nissl staining (right) in the cerebral cortex of mock- and SARS-CoV-2-infected mice at 6 and 15 dpi. Red dashed boxes indicate high-magnification views of the regions with reduced NeuN immunoreactivity. Black dashed lines demarcate areas with reduced NeuN immunoreactivity. Scale bars, 100 µm (low magnification) and 25 µm (high magnification). Representative images from n=3-5 independent biological replicates are shown. Statistical significance for **b** and **d** was determined using a one-way ANOVA with Dunnett’s multiple comparisons test. The exact P values are indicated; N.D., not detected.

Infected mice showed transient weight loss that was recovered by 7–8 dpi, with survival stabilising at approximately 60% (Extended Data Fig. 2a,b). To define the viral tropism and clearance kinetics, we quantified the viral burden across tissues. At 6 dpi, viral RNA was abundant in peripheral tissues, with high levels in the lung, nasal turbinate, and eye, but was undetectable in the testis, and subsequently declined to near-baseline levels by 60-90 dpi (Extended Data Fig. 2c–f). In contrast, viral RNA in the brain peaked at 6 dpi and remained moderately detectable through 90 dpi, indicating prolonged persistence in the brain relative to that in peripheral tissues (Fig. 1b). Given that SARS-CoV-2 infection induces neuroinflammation in this model^27^, we analysed the cytokine and chemokine levels in the brain tissue. Multiplex cytokine profiling revealed robust induction of pro-inflammatory mediators—, including CCL2, CXCL10, IL-6, and TNF-α—, in the brain at 6 dpi, coinciding with the peak viral load (Fig. 1c; Extended Data Fig. 3). This induction was transient, with most analytes returning toward the baseline by 15–30 dpi; however, IL-1β levels remained modestly elevated through 30 dpi before declining. Despite the resolution of acute cytokine induction, we observed persistent neuronal alterations. Transcriptional analysis of the mature neuronal marker *Rbfox3* (NeuN) in whole-brain tissue showed transient downregulation at 6 dpi, which was recovered by 30 dpi (Fig. 1d). In contrast, immunohistochemical analysis revealed persistent NeuN-depleted regions in the cerebral cortex through at least 60 dpi, but not in the olfactory bulb or hippocampus (Extended Data Fig. 4). Distinct areas of reduced NeuN immunoreactivity—, demarcated by black dashed lines—, emerged acutely at 6–15 dpi (Fig. 1e). High-magnification views of these regions (red dashed boxes) confirmed a profound reduction in NeuN immunoreactivity compared with that in mock controls. However, Nissl staining of the adjacent cortical areas revealed that neuronal cell density remained largely intact, although the neurones often exhibited a shrunken or condensed morphology (Fig. 1e). Interestingly, the distribution of this phenotype was spatially heterogeneous; these "patchy" areas appeared in stochastic, multifocal locations across individual mice rather than targeting a fixed cortical subregion (Extended Data Fig. 4b,c). Together, these data indicate that SARS-CoV-2 infection is associated with sustained, spatially heterogeneous reductions in cortical NeuN immunoreactivity, which persist despite the resolution of acute cytokine induction.

### Hypothalamic hypocretin/orexin signalling is suppressed during SARS-CoV-2 infection

Consistent with the peak whole-brain viral RNA load at 6 dpi (Fig. 1b), we profiled the whole-brain transcriptional responses. Multidimensional scaling (MDS) analysis showed segregation by time point, with 6 dpi separating most prominently from the mock-infected and later samples (Fig. 2a). Differential expression analysis at 6 dpi revealed widespread upregulation of immune and antiviral response genes (Fig. 2b), while Gene Ontology (GO) enrichment of the upregulated transcripts highlighted innate immune categories, including regulation of the innate immune response and defence response to the virus (Fig. 2c, top). Canonical interferon-stimulated and inflammatory genes, including *Isg15*, *Cxcl10*, *Ccl5,* and *Irf7*, were among the most strongly expressed transcripts (Fig. 2b). In parallel, the downregulated transcripts were enriched for neuronal signalling-related terms, including the neuropeptide signalling pathway and neurotransmitter receptor activity (Fig. 2c, bottom), indicating impairment of synaptic transmission. Notably, *Hcrt* (hypocretin/orexin) was the top downregulated transcript in the neuropeptide signalling category (Fig. 2d). Network analysis revealed that *Hcrt* functioned as a central node within the suppressed neuropeptide module, exhibiting functional connectivity with *Ucn*, *Avp* and *Sstr4* (Fig. 2e). The concurrent downregulation of Hcrt with stress-regulatory (*Ucn*), neuroendocrine (*Avp*), and sleep-modulating (*Npvf*) factors suggested the a coordinated collapse of hypothalamic homeostatic circuits. Pairwise comparisons across time points indicated that the 6 dpi immune transcriptional program was time-restricted, with most genes returning to mock-like levels by 15 dpi, whereas *Hcrt* expression showed a trend toward recovery throughout the post-acute phase (Extended Data Fig. 5).

**Fig. 2.**
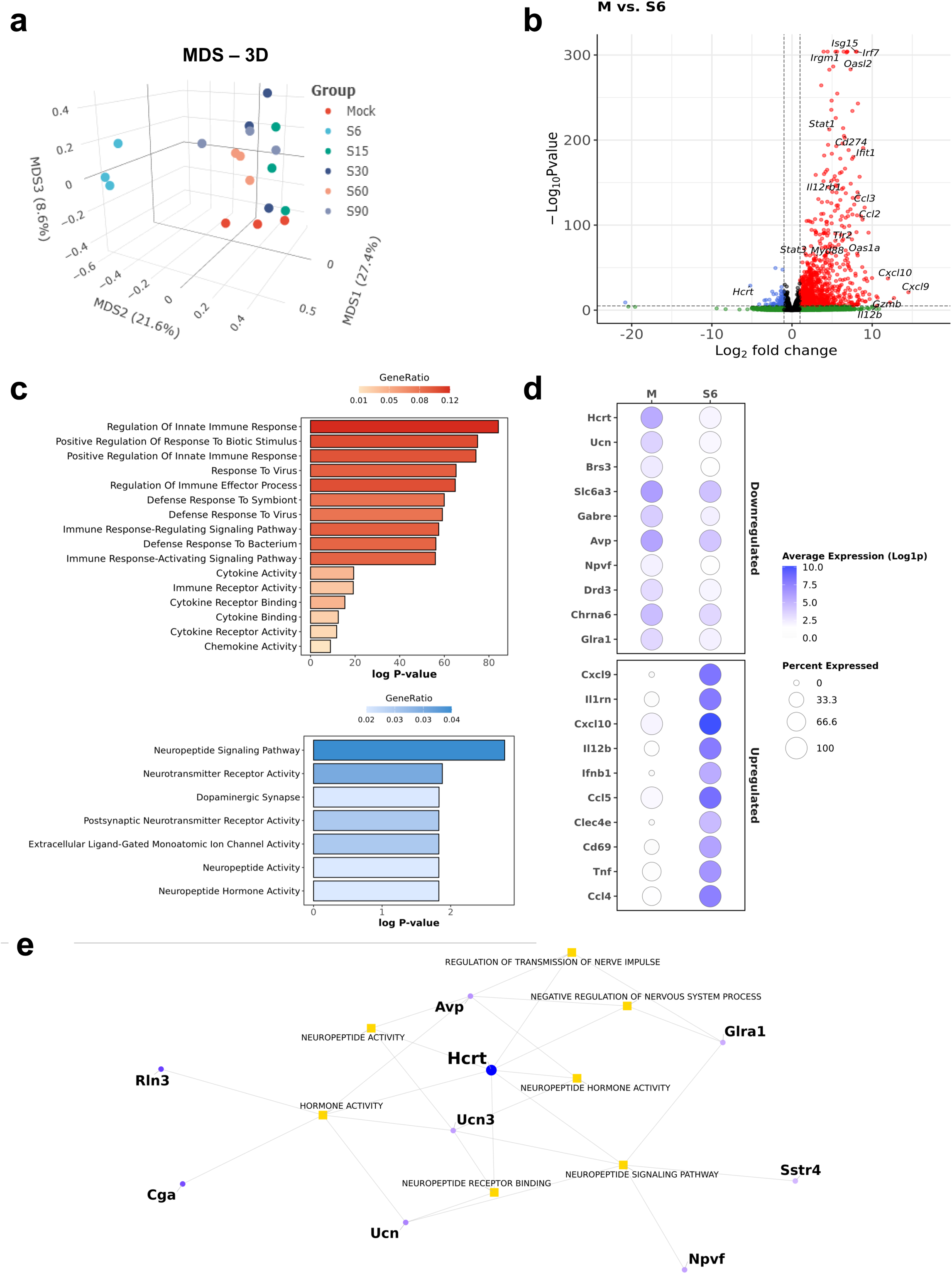
Transcriptomic profiling of SARS-CoV-2-infected K18-hACE2 mouse brains using bulk RNA-seq. **a**, Three-dimensional multidimensional scaling (MDS) plot showing segregation of bulk RNA-seq profiles from mock- and SARS-CoV-2-infected brain samples across time points (Mock, 6, 15, 30, 60, and 90 days post-infection (dpi)); percentages on axes indicate variance explained. **b,** Volcano plot of differentially expressed genes (DEGs) comparing expression at 6 dpi to the mock controls (M vs S6). Vertical dashed lines indicate the log_2_ fold change threshold (|log_2_FC| > 1), and the horizontal dashed line indicates the significance threshold (Benjamini–Hochberg adjusted *P* < 0.01; DESeq2). Genes upregulated at 6 dpi relative to the mock treatment (positive log_2_FC) are shown in red, whereas genes downregulated (negative log_2_FC) at this time point are shown in blue. Selected genes, including *Hcrt* are labelled. **c,** Gene Ontology (GO) enrichment analysis of the upregulated (top) and downregulated (bottom) DEGs at 6 dpi. Bars represent - log_10_(adjusted *P*) of enrichment with bar colour indicating GeneRatio, defined as the proportion of differentially expressed genes annotated to each GO term. **d,** Dot plot showing the expression of selected DEGs with largest absolute log_2_FC across mock and 6 dpi samples, highlighting downregulated neuronal signalling–related genes (top) and upregulated immune-related genes (bottom). Dot size indicates the fraction of samples with detectable expression (non-zero normalised counts) and colour intensity represents the average normalised expression level. **e,** Network representation of the downregulated neuropeptide-associated module, identifying *Hcrt* as a central node connected to additional suppressed targets, including *Ucn*, *Avp* and *Npvf*. Bulk RNA-seq was performed on whole-brain tissues (n = 3 mice per group per time point). Differential expression was assessed using DESeq2 (Wald test) with |log_2_FC| > 1 and Benjamini–Hochberg adjusted *P* < 0.01.

Targeted RT–qPCR and histological analyses were performed to validate these signatures. *Hcrt* mRNA abundance was markedly reduced at 6 dpi and was restored to mock levels by 15 dpi (Fig. 3a). Immunohistochemical staining confirmed that orexin immunoreactivity was restricted to the hypothalamus (Fig. 3b) and profoundly diminished at 6 dpi, and re-emerged from 15 dpi onwards (Fig. 3c). Next, we examined the spatial relationship between the viral antigen and orexin signal using double immunofluorescence staining, which revealed that the orexin signal was minimal within the NP-rich areas, whereas adjacent NP-negative regions retained detectable staining (Fig. 3d, white arrows). Taken together, these data show that acute SARS-CoV-2 infection is associated with the suppression of *Hcrt* transcription and reduced hypothalamic orexin immunoreactivity.

**Fig. 3.**
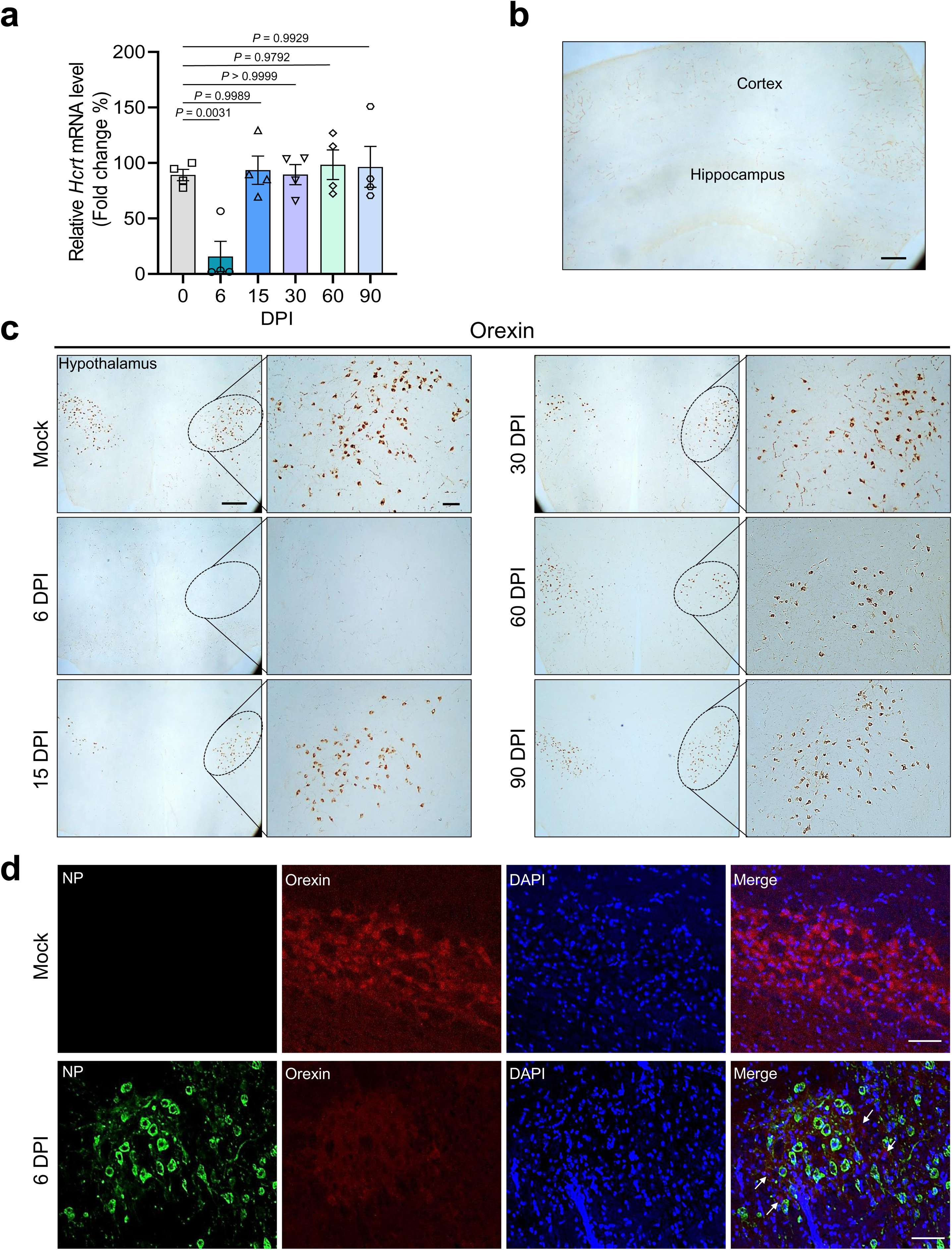
SARS-CoV-2 infection induces transient suppression of hypothalamic hypocretin (orexin) mRNA and protein expression in K18-hACE2 mice. **a**, Time-course quantification of *Hcrt* mRNA levels in whole-brain lysates by RT–qPCR, expressed as fold change relative to mock controls (n = 4 mice per group). **b,** Representative low-magnification immunohistochemistry (IHC) image of orexin staining in a mock-infected brain, showing expression restricted to the hypothalamic region. Scale bar, 100 µm. **c,** Representative IHC images of orexin immunoreactivity in the lateral hypothalamus at the indicated time points. Dashed ovals highlight the orexin neuron field. High-magnification insets (right) show individual neuronal morphology. Scale bars, 100 µm (low magnification), 30 µm (high magnification). **d,** Double immunofluorescence staining for SARS-CoV-2 nucleocapsid (NP; green) and orexin (red) in the hypothalamus in mice infected with SARS-CoV-2 (2 × 10^4^ PFU) at 6 days post-infection (dpi). Nuclei were counterstained with DAPI (blue). White arrows indicate orexin-positive neurones in the NP-negative regions. Scale bar, 25 µm. Representative images in **b-d** from n=3-5 independent biological replicates are shown. Data represent the mean ± s.e.m. Statistical significance was determined using one-way ANOVA with Dunnett’s multiple comparisons test. The exact *P* values are indicated.

### Brain viral burden associates with downregulation of hypothalamic *Hcrt* expression

To determine whether *Hcrt* downregulation is associated with the viral burden in the brain, we quantified the viral RNA and *Hcrt* mRNA across SARS-CoV-2 variants. Upon reanalysing brain tissue from our cohort infected with ancestral (Wuhan-Hu-1) and Beta (B.1.351) SARS-CoV-2 variants^28^, we observed that brain viral RNA increased between 4 and 6 dpi (Fig. 4a), coincident with a progressive decline in *Hcrt* mRNA. In both groups, *Hcrt* expression declined to ∼15–20% of the mock expression at 4 dpi and to below 2% by 6 dpi (Fig. 4b). Infection with the Omicron sublineage KP.3 likewise resulted in a robust brain viral burden at 6 dpi, with *Hcrt* expression at approximately 25% of the mock levels, indicating that *Hcrt* mRNA suppression is conserved across variants (Fig. 4c,d). Next, we investigated whether preventing viral replication through vaccination could rescue *Hcrt* expression. A recombinant RBD subunit protein formulated with alum was administered intramuscularly twice at 2-week intervals, followed by a SARS-CoV-2 challenge (Fig. 4e). Vaccinated mice maintained stable body weight and showed markedly improved survival rates (88%) than the PBS controls (14%) (Fig. 4f,g). At 6 dpi, vaccination reduced the viral RNA levels in the brain by ∼10^4^-fold (Fig. 4h). Notably, *Hcrt* mRNA levels remained near the mock levels (∼90%) in vaccinated mice, rather than decreasing to 3.5%, as in PBS controls (Fig. 4i). Across pooled datasets, brain viral RNA exhibited a strong inverse association with *Hcrt* mRNA abundance (Pearson r = −0.8336; Fig. 4j). Together, these data indicate that suppression of hypothalamic *Hcrt* mRNA inversely associates with brain viral RNA and that vaccine-mediated reductions in brain viral RNA coincide with preservation of *Hcrt* expression.

**Fig. 4.**
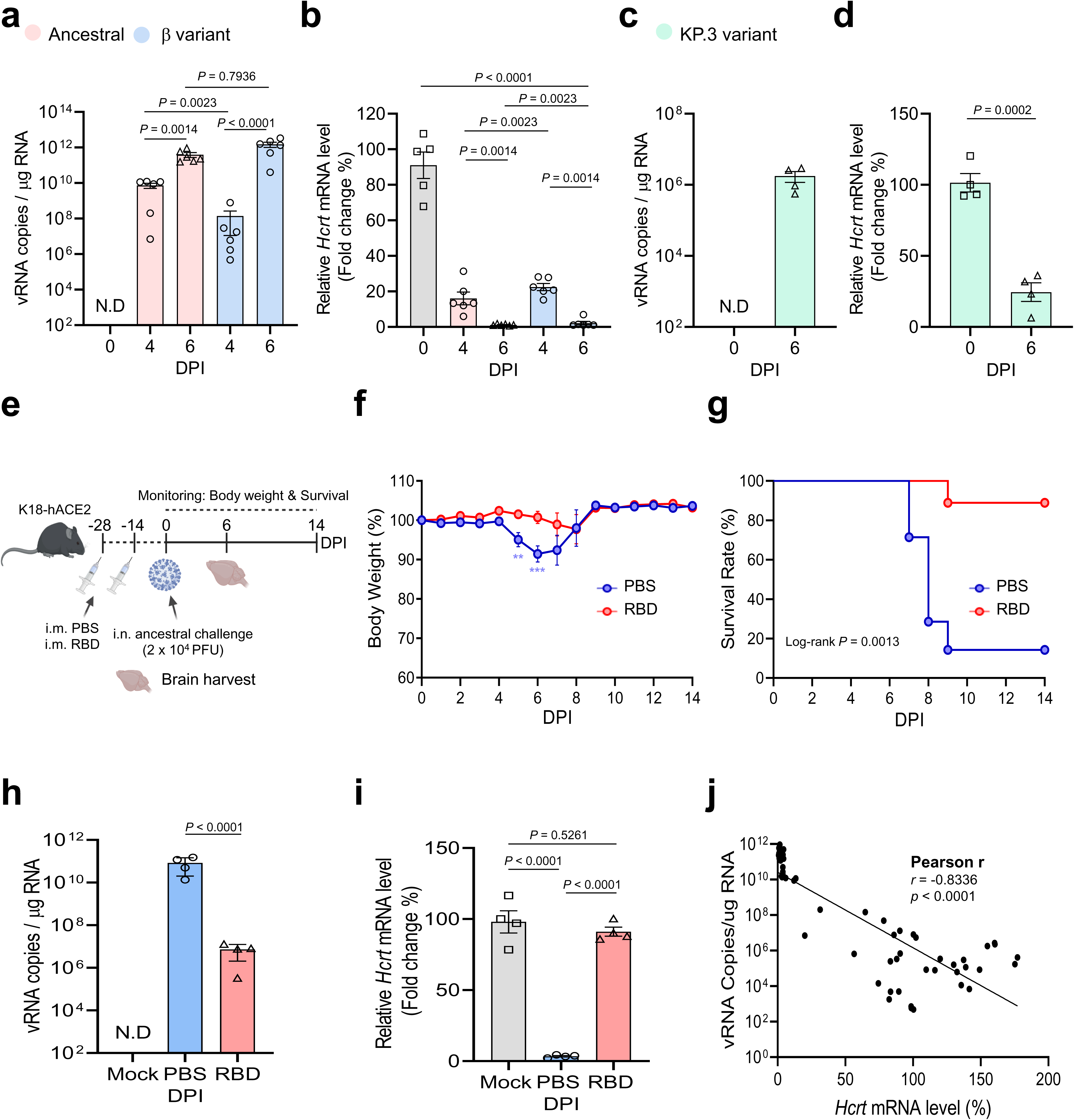
Inverse correlation between SARS-CoV-2 viral burden and transcriptome level of *Hcrt* in K18-hACE2 mice. **a,b**, Quantification of brain viral RNA load (**a**) and relative *Hcrt* mRNA levels (**b**) at 0, 4, and 6 day post-infection (dpi) following infection with ancestral or beta SARS-CoV-2 variants (2 × 10^3^ PFU) (n = 5–6 mice per group). **c,d,** Quantification of brain viral RNA (**c**) and *Hcrt* mRNA levels (**d**) at 6 dpi following infection with the Omicron KP.3 sublineage (n = 4 mice per group). **e,** Schematic representation of the vaccination challenge experiment. K18-hACE2 mice were intramuscularly immunised with two doses of RBD subunit vaccine or PBS and challenged with ancestral SARS-CoV-2 (2 × 10^4^ PFU). **f,g,** Body weight change (**f**, n = 11–13 mice per group) and survival rate (**g**, n = 7–9 mice per group) of vaccinated (RBD) and control (PBS) mice following challenge. For survival analysis, mice were included only if complete follow-up was available through the endpoint, whereas body weight analyses included all animals with longitudinal measurements up to censoring. **h,i,** Quantification of brain viral RNA load (**h**) and *Hcrt* mRNA levels (**i**) at 6 dpi in vaccinated and control mice (n = 4 mice per group). **j,** Correlation analysis between brain viral RNA load and *Hcrt* mRNA levels using pooled data from all experimental cohorts (n = 56 paired samples). Pearson’s correlation coefficients (*r*) and *P* values are indicated. Data represent the mean ± s.e.m. Statistical significance was determined using one-way ANOVA with Dunnett’s multiple comparisons test (**a**,**b**,**i**; comparisons were performed within each experiment against the indicated control), two-tailed unpaired Student’s t-test (**c**,**d**,**h**), two-way ANOVA with repeated measures followed by Sidak’s multiple comparisons test (**f**), or log-rank test (**g**). N.D., not detected.

### MA10 SARS-CoV-2, but not influenza A virus, recapitulates cortical NeuN loss and hypothalamic orexin depletion in wild-type mice

To validate physiological conditions, we extended our analysis to mouse-adapted SARS-CoV-2 (MA10) in wild-type BALB/c mice, which recapitulated the key clinical and pathological features of COVID-19^29,30^. The infected mice exhibited transient weight loss (∼88% at 5 dpi), followed by rapid recovery (Extended Data Fig. 6b). Consistent with previous reports^29^, infection with 4 × 10^5^ PFU led to a high survival rate (approximately 80%), enabling long-term monitoring (Extended Data Fig. 6c).

Next we, quantified the viral burden to assess neuroinvasion dynamics. The viral RNA levels in the lung peaked at 1 dpi and declined thereafter (Extended Data Fig. 6d). Viral RNA levels in the brain peaked at 3 dpi and then gradually decreased (Fig. 5a). RT-qPCR revealed a ∼30% reduction in cortical *Rbfox3* (NeuN) mRNA expression at 6 dpi (Fig. 5b). Beyond this transcriptional deficit, immunohistochemical analysis revealed persistent NeuN-depleted regions in the cerebral cortex up to 90 dpi, whereas the olfactory bulb and hippocampus remained unaffected (Extended Data Fig. 7). Discrete, patchy cortical regions of reduced NeuN immunoreactivity emerged stochastically, with high-magnification images confirming marked NeuN loss despite preserved neuronal density but frequent shrunken morphology on Nissl staining (Fig. 5c; Extended Data Fig. 7a,b).

**Fig. 5.**
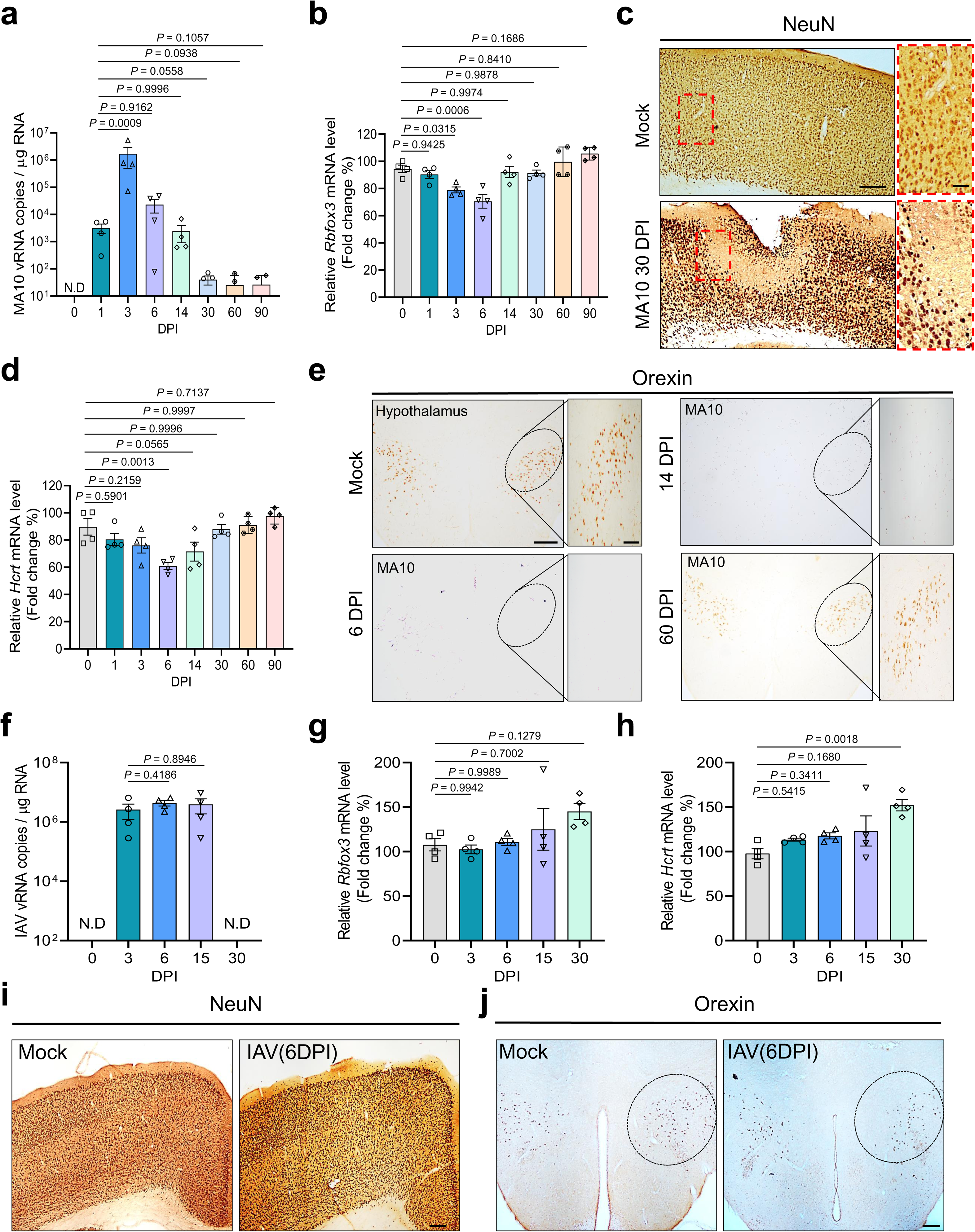
MA10 SARS-CoV-2, but not influenza A virus, induces focal reductions in cortical NeuN immunoreactivity and suppresses hypothalamic orexin in wild-type mice. **a,b**, Time-course quantification of the viral RNA burden (a) and *Rbfox3* (NeuN) mRNA levels (b) in whole-brain lysates of BALB/c mice infected with MA10 SARS-CoV-2 (4 × 10^5^ PFU) (n = 4 mice per group). **c,** Representative NeuN immunohistochemistry (IHC) results in the cerebral cortex of mock- and MA10-infected mice at 30 days post-infection (dpi). Red dashed boxes indicate regions with reduced NeuN immunoreactivity. Scale bars, 100 µm. Representative images from n=3-4 independent biological replicates are shown. **d,** Temporal profile of *Hcrt* mRNA expression in MA10-infected brains (n = 4 mice per group). **e,** Representative orexin IHC in the lateral hypothalamus of mock- and MA10-infected BALB/c mice at 6, 14, and 60 dpi. Dashed ovals highlight the orexin neuron field. Higher-magnification views show orexin-positive neurones. Scale bars, as indicated. Representative images from n=3-4 independent biological replicates are shown. **f–h,** Comparative analysis of C57BL/6 mice infected intranasally with influenza A virus (IAV; PR8 strain, 2 × 10^2^ PFU). Quantification of brain viral RNA (**f**), *Rbfox3* mRNA (**g**), and *Hcrt* mRNA (**h**) levels at the indicated time points (n = 4 mice per group). **i,j,** Representative IHC staining for NeuN in the cortex (**i**) and for orexin in the hypothalamus (**j**) of mock- and IAV-infected mice at 6 dpi. Scale bars, 100 µm. Representative images from n=3-4 independent biological replicates are shown. Data in **a**,**b**,**d**,**f**–**h** represent mean ± s.e.m. Statistical significance was determined using one-way ANOVA with Dunnett’s multiple comparisons test. The exact *P* values are indicated. N.D., not detected.

Tarketed RT-qPCR investigate hypothalamic integrity revealed pronounced depletion of *Hcrt* transcripts (∼40% reduction at 6 dpi), which persisted at 14 dpi (Fig. 5d). Immunohistochemistry confirmed that the reduced orexin expression was restricted to the lateral hypothalamus (Fig. 5e). These findings demonstrate that MA10 infection in wild-type mice reproduces the distinctive cortical and hypothalamic neuropathology observed in K18-hACE2 mice.

To determine whether these lesions are specific to SARS-CoV-2 or represent a generalised consequence of severe respiratory viral infection, we compared these outcomes with those in C57BL/6 mice infected intranasally with influenza A virus (IAV; H1N1 strain PR8). IAV infection led to progressive morbidity, with body weight decreasing to ∼80% by 9 dpi and survival reduced to ∼45% (Extended Data Fig. 8a–c). Despite robust pulmonary replication (Extended Data Fig. 8d) and brain viral RNA levels equivalent to those with MA10 infection (Fig. 5f), transcriptional analysis revealed no significant changes in *Rbfox3* mRNA (Fig. 5g) and NeuN immunoreactivity (Fig. 5i). Furthermore, *Hcrt* mRNA (Fig. 5h) and orexin expression levels remained unaltered (Fig. 5j).

Together, these data reveal that, although MA10 SARS-CoV-2 infection induces prominent cortical neuronal dysfunction and hypocretin depletion, IAV infection does not, despite comparable neuroinvasion. These results indicate that selective suppression of the hypocretin system and cortical neuronal damage represent a distinctive neuropathological signature intrinsic to SARS-CoV-2.

### Hypocretin/orexin signalling restores NeuN expression and neuronal integrity during SARS-CoV-2 infection

We investigated whether impaired orexin signalling could act as a precipitating factor to explore the mechanistic link between hypothalamic dysfunction and cortical neuronal deficits. Human iPSC-derived glutamatergic neurones, which lack endogenous orexin signalling under standard culture conditions^31^, were productively infected with SARS-CoV-2. Although they showed intracellular viral RNA accumulation and robust Nucleocapsid (NP) production, NeuN expression was not reduced (Extended Data Fig. 9). This indicated that the viral burden alone did not account for NeuN downregulation in cortical neurones and instead implicated extrinsic systemic influences, such as disrupted orexin signalling. Next, we examined whether orexin directly modulated NeuN expression (Fig. 6a). Treatment with recombinant orexin-A/B (γORX-A/B) increased NeuN abundance by ∼1.7-fold compared with that in the mock controls, whereas pretreatment with the dual orexin receptor antagonist suvorexant abolished this effect (Fig. 6b), indicating receptor-dependent NeuN upregulation. Dose-response analysis revealed that 10 nM and 100 nM γORX-A/B increased NeuN levels by ∼2.3-fold and ∼2.8-fold, respectively (Fig. 6c). To validate this mechanism *in vivo*, BALB/c mice were infected with MA10 SARS-CoV-2 and treated intranasally with γORX-A/B daily from 0 to 5 dpi (Fig. 6d). At 6 dpi, the brain viral loads were comparable between the vehicle- and γORX-treated groups (Fig. 6e), indicating that orexin administration does not alter viral replication. *Hcrt* mRNA levels were reduced by approximately 40% in both groups (Fig. 6f), consistent with a sustained endogenous orexin deficit, whereas total ORX-A/B levels were significantly higher in γORX-treated brains (Fig. 6g), indicating efficient peptide delivery. Subsequently, western blot analysis of whole brain lysates revealed that NeuN levels in vehicle-treated mice were comparable with those in mock-infected controls, whereas γORX-A/B treatment induced a marked increase in NeuN levels (Fig. 6h). Quantification revealed that NeuN protein abundance was approximately 1.8-fold higher in γORX-A/B-treated mice than in mock controls (Fig. 6i), confirming the robust induction observed *in vitro* (Fig. 6b). Collectively, these data show that exogenous orexin supplementation restores NeuN expression *in vivo* and identify orexin signalling as a potential regulator of neuronal integrity during SARS-CoV-2 infection persisting to Long COVID.

**Fig. 6.**
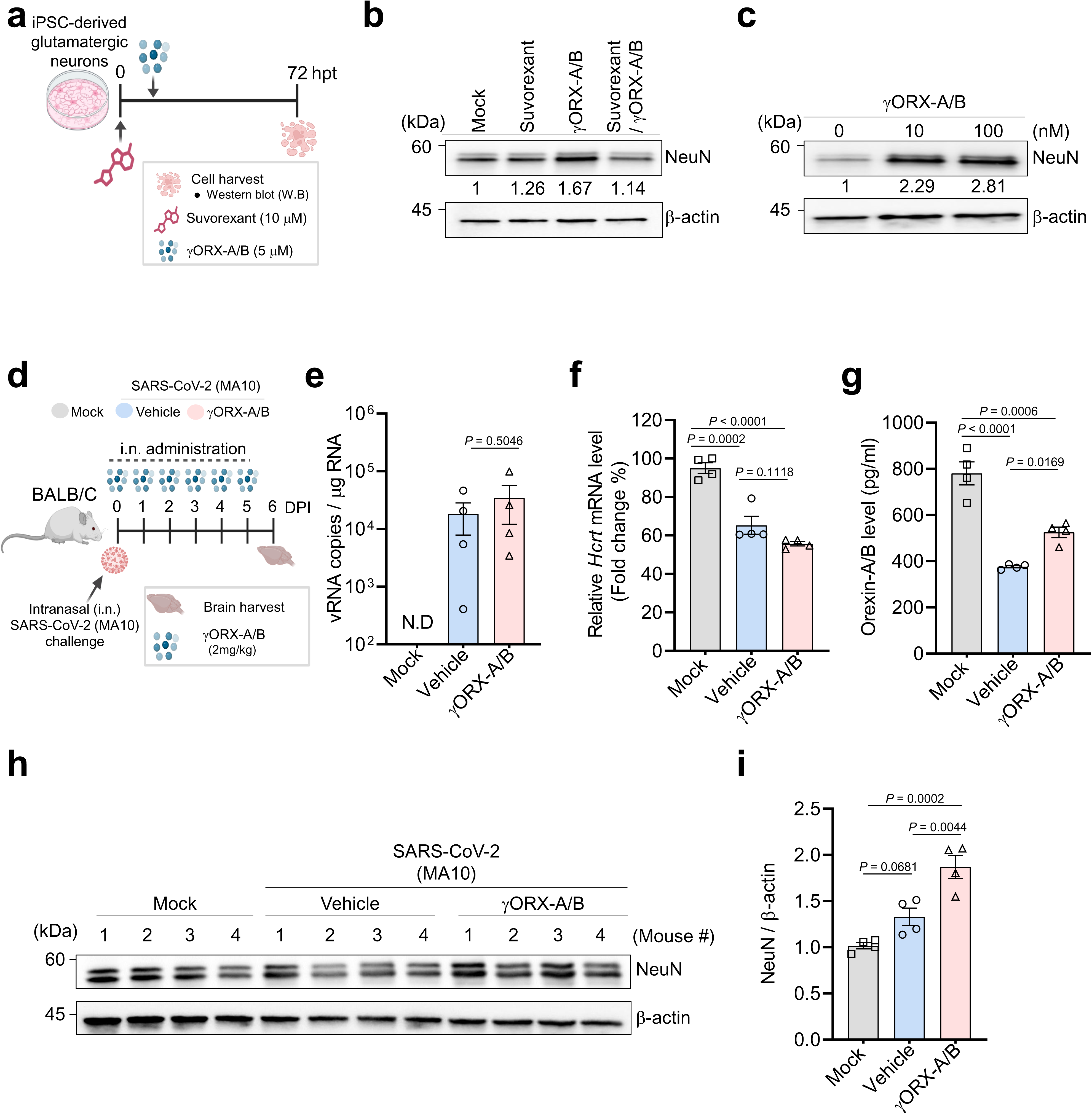
Exogenous orexin signalling promotes NeuN expression *in vitro* and restores cortical neuronal integrity *in vivo* during SARS-CoV-2 infection. **a**, Schematic illustration of the *in vitro* experimental design used to assess the specificity of orexin signalling. Human iPSC-derived glutamatergic neurones were pre-treated with the dual orexin receptor antagonist suvorexant (10 µM) or vehicle for 2 h, followed by treatment with recombinant orexin-A/B peptides (γORX-A/B; 5 μM each) at the indicated concentrations; the cells were then harvested at 72 h post-treatment (hpt) for immunoblotting. **b,** Western blot analysis showing the effect of suvorexant pretreatment on γORX-A/B-mediated NeuN induction. Representative blots (top) and quantification of NeuN protein levels normalized to β-actin (bottom) are shown. The fold change relative to the mock control is indicated below the blots. **c,** Dose-dependent effects of orexin on NeuN expression. Neurones were treated with increasing concentrations of γORX-A/B (0, 10, and 100 nM) and analysed by western blotting at 72 h post-treatment (hpt). **d,** Schematic representation of the *in vivo* rescue experiments. BALB/c mice were intranasally infected with MA10 SARS-CoV-2 (4 × 10^4^ PFU) and administered γORX-A/B (2 mg kg ¹) or vehicle daily from 0 to 5 days post-infection (dpi). Brain tissues were collected at 6 dpi. **e,f,** Quantification of viral burden (**e**) and *Hcrt* mRNA levels (**f**) in whole-brain lysates at 6 dpi using RT–qPCR. **g,** ELISA quantification of the orexin-A/B peptide concentrations in brain homogenates. **h,** Representative Western blots of NeuN expression in whole-brain lysates from mock, vehicle-treated and γORX-A/B-treated mice at 6 dpi. **i,** Densitometric quantification of NeuN protein levels normalised to β-actin. Data in **e**–**g** and **i** represent mean ± s.e.m. (n = 4 mice per group). Statistical significance was determined using one-way ANOVA with Dunnett’s multiple comparison test (**f**,**g**,**i**) or a two-tailed unpaired Student’s t-test (**e**). The exact *P* values are indicated. N.D., not detected.

## Discussion

SARS-CoV-2 is increasingly recognised as a cause of persistent neurological sequelae, yet the mechanisms linking acute infection with long-term circuit dysfunction remain unclear. We identified two convergent *in vivo* signatures of SARS-CoV-2 neuropathology: focal, heterogeneous loss of cortical NeuN immunoreactivity persisting beyond the acute phase, and selective suppression of hypothalamic hypocretin/orexin (*Hcrt*) expression during early infection. Using comparative viral models and pharmacological interventions, we revealed that these processes are connected through orexinergic circuit disruption, suggesting a pathway that bridges acute infections to lasting neuronal vulnerability. This mechanism provides a potential cellular basis for Long COVID symptoms^32^, including sleep–wake cycle instability, fatigue, and cognitive dysfunction.

Neuroinflammation is a hallmark of respiratory viral infection and can profoundly influence neuronal function; however, whether inflammation alone explains persistent, spatially restricted neuronal dysfunction remains uncertain. In our analysis, inflammatory mediators such as Cxcl10 and Ccl2 surged during the acute phase (Fig. 1c), consistent with reports of blood–brain barrier disruption and cytokine induction^33,34^—, but subsided thereafter, whereas the downregulation of focal cortical NeuN persisted. This temporal dissociation indicates that early cytokine elevation cannot fully account for the enduring and anatomically focal NeuN downregulation (Extended Data Fig.4). Comparative viral evidence reinforces the “inflammation-is-not-enough” model: peripheral H1N1 influenza infection triggers robust neuroimmune activation, but does not produce the observed shrinkage or NeuN loss in neurones^13,14^. Thus, the mammalian brain can withstand substantial inflammatory stress without neurodegeneration, implying that persistent cortical pathology requires mechanisms beyond general neuroinflammatory signalling. This is consistent with recent CSF profiling in Myalgic Encephalomyelitis/Chronic Fatigue Syndrome (ME/CFS), which suggests that distinct central immune signatures, rather than generalised inflammation alone, drive specific neurological phenotypes^35–38^.

We tracked the mature neuronal marker NeuN (*Rbfox3*) and observed sustained, spatially “patchy” reduction in cortical NeuN immunoreactivity, appearing as stochastic multifocal clusters. Nissl staining indicated that neurones within NeuN-attenuated foci were not absent but often appeared shrunken or atrophic. Importantly, loss of NeuN antigenicity does not necessarily indicate neuronal death but often reflects metabolic stress or transcriptional suppression in structurally preserved neurones, as shown in ischaemic models^39–41^. Nonetheless, persistent NeuN attenuation, indicates a chronically compromised neuronal state that predisposes patients to delayed neuronal degeneration or dysfunction^42^. Thus, following SARS-CoV-2 infection, neurones survive but adopt a shrunken, transcriptionally altered state, which may underlie the circuit instability and cognitive ’brain fog’ associated with Long COVID.

The next question was whether these neuropathological features depend on direct viral access to the CNS. Our data identified SARS-CoV-2 as a neurotropic virus capable of entering the CNS and exhibiting tropism for hypothalamic circuits. High-resolution imaging at 6 dpi revealed an inverse spatial relationship between viral burden and orexin expression: orexin immunoreactivity was virtually absent in regions dense with viral NP signals, whereas adjacent NP-negative zones retained detectable staining (Fig. 3d, white arrows). This spatial exclusivity suggests that orexin loss arises from local viral infection rather than systemic inflammation. Human autopsies and animal studies likewise confirm SARS-CoV-2 neuroinvasion^10,11^. Similar patterns are found in neuroadapted influenza strains; for instance, neurotropic H1N1 invades via the olfactory route to destroy orexin neurones and induce narcolepsy-like sleep disruption^43^ and H7N7 infection drives microglial activation and synaptic loss^44^. In contrast, our PR8 (H1N1) model—, a well-characterised non-neurotropic strain that induces severe systemic inflammation but lacks intrinsic neuronal replicative capacity^45,46^—, achieved comparable brain viral RNA loads but failed to elicit hypothalamic orexin dysfunction or NeuN attenuation. This dissociation demonstrates that systemic inflammatory severity and the mere presence of viral RNA in the brain are insufficient to drive this pathology; rather, it indicates that disruption of the orexin-NeuN axis reflects a specific neuropathological signature intrinsic to SARS-CoV-2 infection in the hypothalamus.

We thus propose a unified framework wherein selective hypothalamic orexin suppression is the primary driver of cortical neuronal dysfunction. Our transcriptomic analysis revealed synchronised suppression of *Hcrt* along with *Ucn*, *Avp* and *Npvf*, consistent with a coordinated perturbation of hypothalamic homeostatic programs. Given that CRF/urocortin-family signalling and vasopressin directly modulate orexin neuronal activity and orexin-dependent control of stress-associated arousal and wakefulness^47,48^, and that Neuropeptide VF (Npvf) exhibits evolutionarily conserved links to orexin cell states and behavioral-state regulation^49,50^, their concurrent downregulation implies a systemic failure of the arousal network. Orexin neurones serve as integrative hubs that regulate sleep—wake cycle stability, arousal, and neuroendocrine balance, while maintaining neuroprotection and immune homeostasis^51^. Orexin-A promotes neuronal survival under oxidative stress^20^, supports hippocampal neurogenesis^52^ and limits neuroinflammation via NF-κB inhibition^21^. This central role aligns with the emerging evidence of orexinergic disruption in COVID-19, including autoantibodies against orexin receptors^53^. To separate the direct viral effects from circuit-level mechanisms, we used human iPSC-derived cortical neurones, which supported viral replication but maintained NeuN expression without orexin signalling (Extended Data Fig. 9), indicating that direct infection alone was insufficient for NeuN loss. Conversely, exogenous orexin restored the cortical NeuN levels in a receptor-dependent manner without altering the viral burden (Fig. 6i). Overall, these data suggest that virus-mediated disruption of the orexin system offers a plausible basis for the fatigue, arousal deficits, and cognitive dysfunction characteristic of Long COVID^32^.

The K18-hACE2 transgenic mouse is a standard model for severe COVID-19 and has been used to assess vaccine efficacy and respiratory disease pathology^34,54^. Beyond lung injury, it recapitulates extrapulmonary symptoms, including retinal inflammation and ocular discharge, which mirror conjunctivitis in patients^55,56^. These manifestations align with known neuronal dissemination pathways connecting the respiratory, neural, and ocular tissues, supporting their use in studying SARS-CoV-2 neuroinvasion^10,56^. In this model, infection reduced hypothalamic orexin and cortical NeuN expression, a phenotype conserved across variants including Beta and Omicron KP.3 (Fig. 4a-d), indicating that orexin dysfunction is an intrinsic feature of SARS-CoV-2 neuropathology. To confirm its physiological relevance, we analysed MA10 SARS-CoV-2 infection in wild-type BALB/C mice^57^, which reproduced the same deficits. Unlike K18-hACE2 mice, which showed partial orexin recovery by 15 dpi, MA10-infected mice showed prolonged *Hcrt* suppression through 14 dpi and persistent cortical NeuN loss up to 90 dpi. This sustained pathology in wild-type mice demonstrates that SARS-CoV-2-induced neuropathology is a robust and intrinsic outcome, and not an artefact of transgenic ACE2 expression.

This study has some limitations that should be considered for future studies. First, NeuN attenuation alone does not indicate neuronal death. To determine whether this reduction in NeuN immunoreactivity signifies irreversible neurodegeneration or a reversible functional state, multi-marker analyses using Fluoro-Jade, activated caspase-3, and synaptic markers within NeuN-low regions are required at extended time points following viral clearance^39^. Second, whether orexin deficiency is causal or correlative remains unresolved. Although exogenous orexin restores NeuN levels, targeted loss-of-function perturbations of orexin neurones or receptors combined with longitudinal behavioural profiling are required to establish causality^58^. As the constraints of biosafety level (BSL)-3 limit behavioural assays, future studies integrating telemetric sleep–wake recording and cognitive testing are needed for linking molecular changes to the Long COVID-relevant functional deficits. Furthermore, unlike hypothalamic orexin neurones, the iPSC-derived glutamatergic neurones used in this study do not secrete orexin, therefore, extending the mechanistic analyses to orexinergic neuronal models will clarify whether orexin-producing cells themselves are directly vulnerable to infection.

Collectively, our findings identify hypocretin/orexin suppression as a selective feature of SARS-CoV-2 brain pathology, which coincides with persistent, spatially heterogeneous cortical NeuN attenuation. Restoring orexin signalling rescued NeuN expression without altering the viral load, supporting the idea that disrupting the orexinergic circuitry may lead to neuronal dysfunction and contribute to the arousal and cognitive impairments characteristic of Long COVID.

## Methods

### Ethics statement and biosafety

All animal procedures were approved by the Institutional Animal Care and Use Committee (IACUC) of the Korea Research Institute of Chemical Technology (KRICT) and conducted in accordance with the institutional guidelines and relevant national regulations (Protocol ID 8A-M6; IACUC IDs 2023-8A-07-01, 2023-8A-08-02, 2025-8A-04-03 and 2025-8A-08-02).

Experiments involving infectious SARS-CoV-2 were performed in biosafety level 3 (BSL-3) facilities at KRICT, and experiments involving influenza A virus (IAV; H1N1) were performed in biosafety level 2 (BSL-2) laboratories. All studies were performed in compliance with the relevant national and institutional biosafety regulations.

### Animals and in vivo procedures

Male B6.Cg-Tg(K18-hACE2)2Prlmn/J mice were purchased from The Jackson Laboratory (Bar Harbor, ME, USA). Male BALB/cAnNCrlOri and C57BL/6NCrlOri mice were obtained from Orient Bio Inc. (Seongnam, Republic of Korea). Animals were maintained in a BSL-2 animal facility in individually ventilated cages at the Korea Research Institute of Chemical Technology (KRICT), with controlled temperature (22 ± 2 °C), humidity (50 ± 10%), and a 12-h light/dark cycle. The experimental groups comprised age-matched mice (8–9 weeks old).

All viral inoculations [SARS-CoV-2 (50 PFU), MA10 SARS-CoV-2 (4 × 10 PFU), and influenza A virus (2 × 10² PFU)] were administered intranasally (IN) at a volume of 20 μL per mouse, under isoflurane anaesthesia in BSL-3 or BSL-2 animal facilities, as appropriate, and all efforts were made to minimise animal suffering. IN inoculations were performed as previously described^56^. Mock-infected mice were administered an equal volume of PBS. Post-infection, body weight and survival were monitored daily. At designated time points following infection (as indicated in the respective figure legends), mice were deeply anaesthetized with isoflurane and perfused transcardially with PBS, followed by 4% paraformaldehyde (PFA) for histological assessment, or fresh tissues were harvested and flash-frozen for downstream molecular and biochemical analyses.

To evaluate vaccine efficacy, K18-hACE2 mice were immunised intramuscularly (IM) with 10 µg of recombinant SARS-CoV-2 Spike RBD protein formulated using aluminium hydroxide gel (Alum) (Alhydrogel 2%, InvivoGen, San Diego, CA, USA) as an adjuvant in a 1:1 ratio, at a final volume of 100μL per mouse or with PBS/Alum (Mock control). Booster immunisations with the same dose and formulation were administered after two weeks. Two weeks after the final vaccination, mice were challenged intranasally with 2 × 10^4^ PFU of SARS-CoV-2 in a 20 μL volume (100MLD_50_). The mice were then monitored daily for 14 dpi. On day 6 post-infection, brain tissues were collected for downstream molecular and biochemical analyses as described above.

To evaluate the therapeutic effects of orexin in vivo, BALB/c mice were intranasally infected with MA10 SARS-CoV-2 (4 × 10^4^ PFU). Recombinant orexins A and B (2 mg/kg each; MCE, HY-106224B, HY-P1349) or vehicle (PBS) were administered intranasally daily from 0 to 5 dpi. Mice were euthanised at 6 dpi and whole brains were harvested for viral load quantification and western blot analysis.

### Cells and viruses

The ancestral SARS-CoV-2 Korean strain (GISAID: EPI_ISL_407193; NCCP: 43326) and the Omicron KP.3 variant (NCCP: 43496) were obtained from the Korea Disease Control and Prevention Agency (KDCA). The mouse-adapted SARS-CoV-2 MA10 strain was obtained from BEI Resources (NIAID, NIH, USA). All viruses were propagated in Vero cells (CCL-81; ATCC) grown in Dulbecco’s Modified Eagle’s Medium (DMEM) supplemented with 2% foetal bovine serum (FBS) as previously described^57,59^. The influenza A virus H1N1 (PR8 strain) was kindly provided by Prof. Heung Kyu Lee (KAIST). Viral titres were determined using a plaque assay on Vero cells.

### Differentiation of human iPSC-derived glutamatergic neurones

Glutamatergic neurones were generated from human iPSCs as previously described^31^ with minor modifications. Human iPSCs (ATCC-DYR0100; ACS-1011) were cultured to ∼70% confluence in mTeSR1 medium supplemented with Y-27632 (10 µM; Millipore Sigma). Cells were dissociated with 0.5 mM EDTA and seeded (3.5 × 10 cells/well) onto Matrigel-coated 6-well plates in chemically defined medium (CDJ; Neurobasal/DMEM-F12 [1:1] supplemented with B27, N2, and penicillin-streptomycin). On the following day, cells were transduced with lentiviral vectors encoding hNGN2 (Addgene 62223) and rtTA (Addgene 20342) at a multiplicity of infection (MOI) of 5 via spin-infection (1000 × g, 1 h). At 24 h post-transduction, medium was replaced with CDM containing doxycycline (2.5 µg/ml) and puromycin (1 µg/ml) for inducing NGN2 expression and selection, respectively. After 48 h of selection, cells were re-seeded on Matrigel-coated plates and maintained in neuronal growth medium (NGM; CDM supplemented with 2.5% FBS, 20 ng/mL BDNF, 20 ng/mL GDNF, 1 µM cAMP, 100 µM ascorbic acid) containing doxycycline (2.5 µg/mL) and the Notch inhibitor DAPT (2.5 µM; Selleckchem), for 5 days. Neuronal maturation continued in NGM without DAPT for an additional week. Finally, neurones were maintained in BrainPhys™ medium supplemented with 2.5% FBS, B27, N2, GlutaMAX, and penicillin-streptomycin before the experiments.

### In vitro infection and pharmacological treatment

Differentiated neurones (5 × 10 cells/well) were seeded into 12-well plates and inoculated with SARS-CoV-2 at a MOI of 1. After 1 h, the inoculum was removed, the cells were washed twice with PBS, and fresh medium was added.

For orexin rescue experiments, recombinant orexin-A and orexin-B (MCE, HY-106224B, HY-P1339B) were added at the indicated concentrations (10 nM, 100 nM, or 5 µM) immediately after infection. To assess receptor specificity, neurones were pre-treated with the dual orexin receptor antagonist Suvorexant (10 µM; Biorbyt, orb146219) or vehicle (DMSO) for 2 h prior to infection and subjected to orexin treatment. The cells were harvested at 72 hours post-infection (hpi) for downstream RNA and protein analyses.

### Quantitative RT-PCR

Brain tissues from K18-hACE2 mice infected with the ancestral SARS-CoV-2 or the Beta variant were generated as described previously by Lee et al^28^. Total RNA was extracted from tissues using the Maxwell RSC Simply RNA Tissue Kit (AS1340; Promega, Madison, WI, USA) according to the manufacturer’s instructions. Quantitative RT-PCR was performed on a QuantStudio 3 Real-Time PCR System (Applied Biosystems, Foster City, CA, USA) using either the One-Step PrimeScript III RT-qPCR Mix (RR600A) or the One Step TB Green PrimeScript RT-PCR Kit II (RR086A) (Takara, Kyoto, Japan), following the manufacturer’s protocols. Viral RNA targeting the SARS-CoV-2 nucleocapsid (N) gene was detected using the 2019-nCoV-N1 probe (10006770; Integrated DNA Technologies, Coralville, IA, USA). The expression of *Hcrt*, *Rbfox3* and β*-actin* genes was analysed using customised primers (Bionics, Korea). Primer sequences used in this study are listed in Table S1. Average threshold cycle (Ct) values of *Hcrt* and *Rbfox3* from the PCRs were normalised to the average Ct values of β*-actin*.

### RNA-seq and analysis

The sequencing library was prepared using the TruSeq Stranded mRNA Sample Prep Kit and sequenced on a NovaSeq 6000 (Illumina, San Diego, CA, USA), yielding more than 6G bases of sequence per sample. Adaptor sequences were removed from the sequenced reads using Trimmomatic (version 0.39)^60^ and the processed reads were aligned to the mouse reference genome (GRCm38/mm10) using HISAT2 (version 2.2.1)^61^. The aligned reads were quantified at the gene level using FeatureCounts (version 2.0.6)^62^. Differential expression analyses were performed using DESeq2 (version 1.42.1)^63^. Differentially expressed genes were visualised as volcano plots generated with the EnhancedVolcano package (version 1.20.0) and identified using the Wald test with abs (log_2_ fold change) > 1 and adjusted P-value (Benjamini–Hochberg) < 0.01 as the cut-off. Multidimensional scaling analysis was performed using the cmdscale function in the R stats package (version 4.3.3) based on genes with non-zero expression and non-zero variance among samples. Gene Ontology (GO) enrichment analysis was conducted for DEGs using the enrichGO function in the clusterProfiler package (version 4.10.1)^64^, with gene annotations from the org.Mm.eg.db database. Enriched GO terms were defined by P-value < 0.05 and q-value < 0.05. Associations between enriched GO terms and DEGs were represented as networks constructed using igraph (version 2.2.1)^65^ and visualised using ggraph (version 2.2.2) in R.

### Multiplex profiling

The brains of SARS-CoV-2-infected mice were dissected at 0, 6, 15, 30, 60, and 90 dpi and homogenised in bead tubes (a-PSBT, GeneReach Biotechnology, Taichung, Taiwan). These homogenates were aliquoted and analysed with the Luminex mouse cytokine/chemokine magnetic bead panel (LXSAMSM-21; LXSAMSM-03, R&D Systems, Minneapolis, MN, USA) using a Bio-Plex 200 multiplexing instrument (Bio-Rad, Hercules, CA, USA) to assess the cytokine/chemokine expression.

### Immunohistochemistry

As previously described^66,67^, mouse brain tissues were coronally prepared at 30 μm thickness from the olfactory bulb to cerebellum. The sections were rinsed in PBS and incubated overnight with mouse anti-NeuN (1:1000; Merck Millipore) and anti-orexin/prepro-orexin (AB3096; Merck Millipore) antibodies for staining general neurones. The next day, the tissue sections were rinsed with 0.5% bovine serum albumin in PBS and incubated for 1 h at room temperature (RT) with a mouse biotinylated secondary antibody (1:400; KPL, MD, USA). This was followed by incubation with an avidin-biotin-peroxidase complex (Vectastain ABC Kit; Vector Laboratories) for visualisation. The bound antiserum was then visualized by incubating with 3,3’ diaminobenzidine (DAB: Sigma) solution under a bright-field microscope (Olympus Optical, Tokyo, Japan).

For Nissl Staining, whole brain sections were mounted on Aminosilane-coated slides and dried for 1 h at RT, dehydrated in 100% ethanol, cleared in xylene, hydrated in a decreasing alcohol gradient, stained with 0.5% cresyl violet (Sigma), washed in distilled water, dehydrated in 100% ethanol, mounted with a coverslip, and viewed under a bright-field microscope (Olympus Optical).

### Immunofluorescence

Immunofluorescence assays were performed as described by Jeong et al^68^. Briefly, mouse brains were perfused and fixed in 4% paraformaldehyde, sectioned at 30 μm, and subjected to standard permeabilisation and blocking procedures. Sections were incubated with primary antibodies, anti-SARS-CoV-2 nucleocapsid protein (40143-MM05, Sino Biological), and anti-orexin/prepro-orexin (AB3096, Merck Millipore), followed by incubation with Alexa Fluor 488-conjugated anti-rabbit antibody (A32731, Thermo Fisher Scientific, Waltham, MA, USA) and Alexa Fluor 594-conjugated anti-mouse antibody (A32744, Thermo Fisher Scientific). Nuclei were counterstained with DAPI. Immunofluorescence was observed using confocal microscopy (LSM700; Carl Zeiss, Oberkochen, Germany).

### ELISA

Orexin levels were quantified in brain homogenate supernatants prepared as described above. The concentration of each was determined using a Mouse Hcrt(Orexin) ELISA Kit (EM0453, FineTest) according to the manufacturer’s instructions.

### Western blotting

For protein analysis, cultured neurones or brain tissue were lysed in radioimmunoprecipitation (RIPA) buffer (Thermo Fisher Scientific), and proteins in the lysate were separated on a denaturing polyacrylamide gel and transferred to a polyvinylidene fluoride (PVDF) membrane (Merck Millipore, Burlington, MA, USA). The membrane was blocked with 5% skimmed milk (BD Biosciences) in Tris-buffered saline with 0.1% Tween 20 (TBST) buffer and then incubated with the primary antibodies anti-SARS-CoV-2 NP (1:1000, 40143-R001, Sino Biological), anti-NeuN (1:1000, A19086, ABclonal), and anti-β-actin (1:5000, sc-47778, Santa Cruz Biotechnology). Horseradish peroxidase (HRP)-conjugated secondary antibodies (Bio-Rad) and enhanced chemiluminescence (ECL) reagents (Thermo Fisher Scientific) were used for protein band detection. Immunoblot band intensities were quantified by densitometry using ImageJ^69^.

### Statistical analysis

Data were analysed using the GraphPad Prism 8.4.3 software (GraphPad Software, San Diego, CA, USA). For multiplex cytokine profiling, values falling below the limit of detection (LOD) were handled based on the analysis type: for heatmap visualisation, undetected values were imputed as 1.0 to facilitate log_2_ transformation (yielding a value of 0). For quantitative comparisons and extended data (Violin plots), these values were assigned as 0 to accurately reflect sample sizes and non-responders. Outliers were identified individually for each analyte and removed using the ROUT method (Q = 1%). Data are presented as the mean ± s.e.m. For comparisons between two groups, an unpaired two-tailed Student’s t-test was used. For comparisons among three or more groups, one-way analysis of variance (ANOVA) with Dunnett’s multiple comparison test was used. Long-term assessment of body weight data was performed using a two-way ANOVA with repeated measures, followed by Sidak’s multiple comparisons test. Survival curves were compared using the log-rank (Mantel–Cox) test. The correlation between viral load and gene expression was assessed using Pearson correlation analysis. A *P-value*< 0.05 was considered statistically significant. The specific statistical tests and sample sizes used are indicated in the figure legends. Schematic overviews for animal experiments were created with BioRender.com.

## Acknowledgement

Graphic images in figures were created using BioRender.

This work was supported by the Korea Research Institute of Chemical Technology (KRICT) under project no. KK2633-20 (A Study on the Next-Generation Infectious Disease Control Technology Program) and by the Korea Institute of Toxicology under project no. KK-2505-01 (2710086913). This research was also supported by the National Research Foundation of Korea (NRF) grant funded by the Ministry of Education, Science, and Technology (MSIT) of the Korean government (RS-2023-00208568 to Y.-C.K.). In addition, this work was supported by the G-LAMP Program of the National Research Foundation of Korea (NRF) grant funded by the Ministry of Education (RS-2025-25441317 to J.-K.R.) and by the National Research Foundation of Korea (NRF) grant funded by the Ministry of Science and ICT (RS-2021-NR061451 to J.-K.R.).

## Author contributions

Conceptualization: G.Y.Y., Y.-C.J., W.-H.S, and Y.-C.K. ; Methodology: G.Y.Y., Y.-C.J. W.- H.S, and Y.-C.K. ; Investigation: G.Y.Y., Y.-C.J., J.-H.C., Y.H., S.Y.S., K.B.K., D.Y.K., and W.Y.H. ; Resources: D.-G.A., J.-K.R, W.-H.S., and Y.-C.K. ; Data curation and analysis: G.Y.Y., Y.-C.J., Y.H., J.-K.R., W.-H.S., and Y.-C.K. ; Writing–original draft: G.Y.Y. and Y.-C.J. ; Writing–review and editing: G.U.J., K.-D.K., J.-K. R., W.-H.S., and Y.-C.K., ; Visualization: G.Y.Y., Y.-C.J., J.-K.R., W.-H.S, and Y.-C.K. ; Supervision : W.-H.S., and Y.-C.K. ; Funding acquisition : Y.-C.J., J.-K.R., W.-H.S., and Y.-C. K.

---

**Extended Data Fig. 1.**
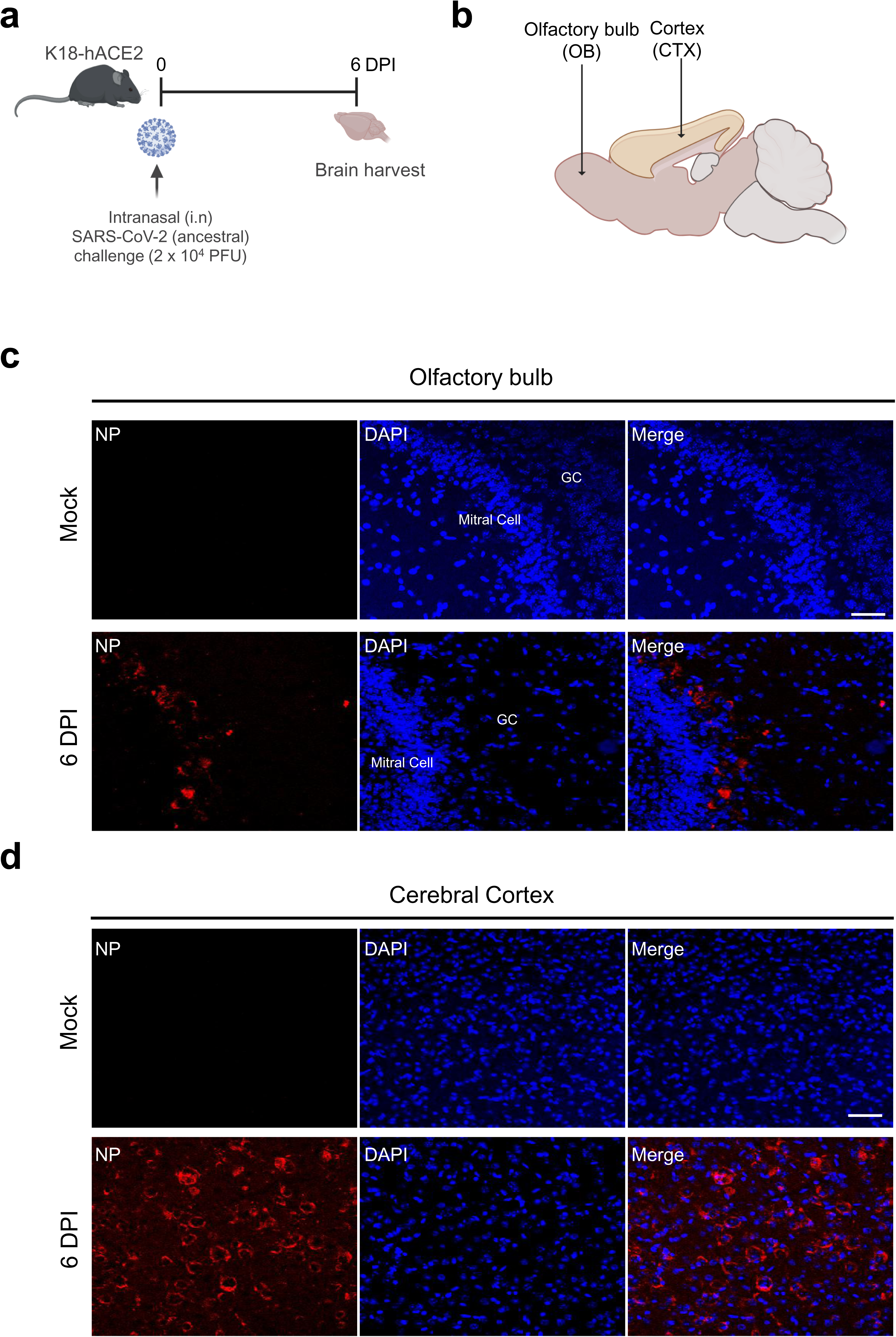
Extensive brain infection and neuroinvasion in K18-hACE2 transgenic mice infected with a lethal dose of SARS-CoV-2. **a,** Schematic of the lethal-dose infection model. K18-hACE2 mice were intranasally challenged with 2 × 10^4^ PFU of SARS-CoV-2 and analysed at 6 dpi. **b,** Diagram of the mouse brain indicating the anatomical regions analysed for viral distribution: olfactory bulb (OB), cortex (CTX). **c,d,** Representative immunofluorescence images of the SARS-CoV-2 nucleocapsid (NP; red) in the olfactory region (**c**) and cerebral cortex (**d**) at 6 dpi. Nuclei are counterstained with DAPI (blue) and merged images are shown. Scale bars, 25 µm. Images are representative of three n = 3 mice.

**Extended Data Fig. 2.**
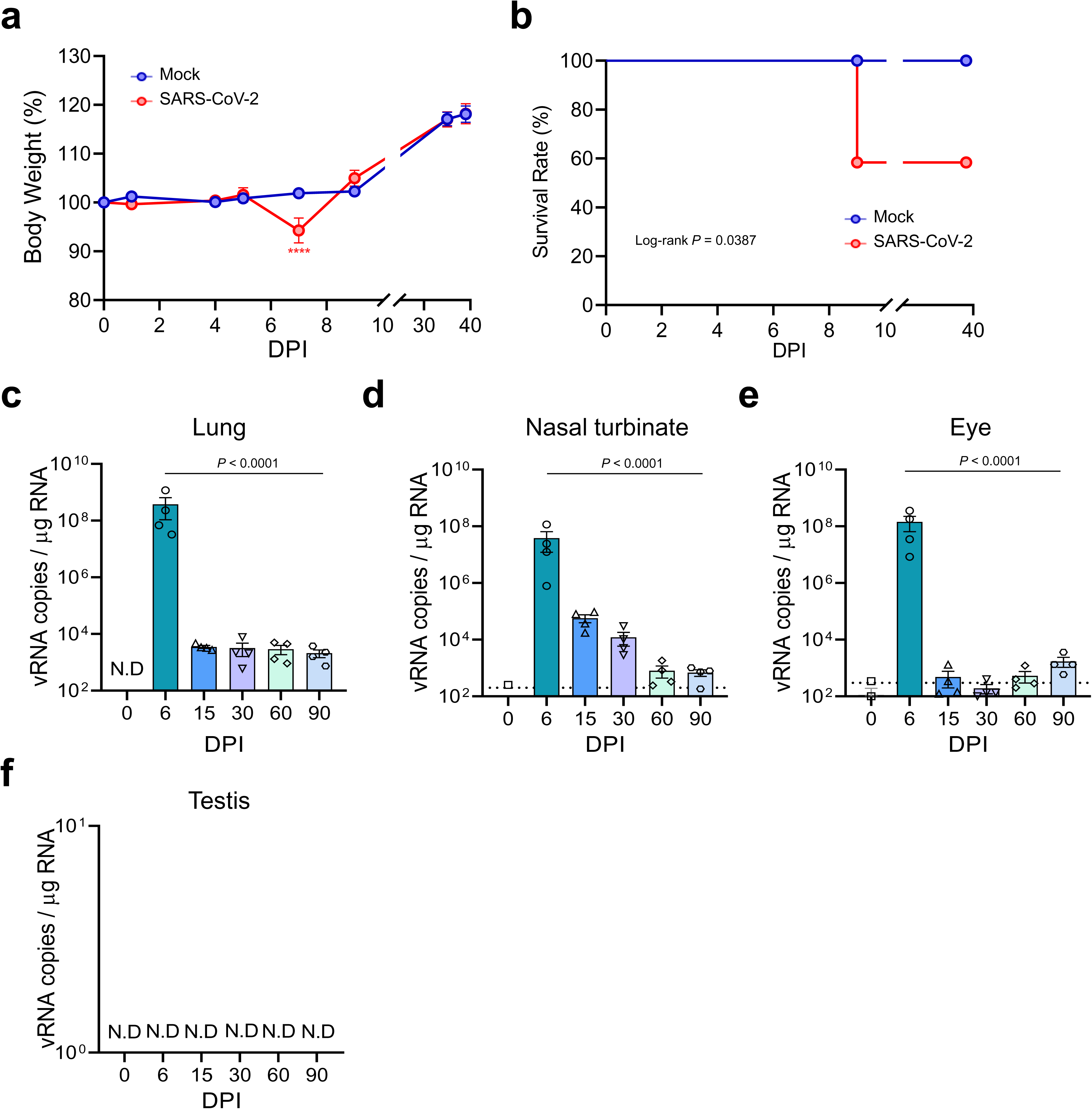
Clinical course and peripheral viral RNA kinetics following SARS-CoV-2 infection in K18-hACE2 mice. **a,b**, Body weight change (**a**; n = 12 mice per group) and survival (**b**; n = 13 mice per group) of K18-hACE2 mice infected intranasally with 50 PFU of SARS-CoV-2 compared with mock controls. **c–f**, Time-course quantification of viral RNA in the lung (**c**), nasal turbinate (**d**), eye (**e**) and testis (**f**) at the indicated time points using RT-qPCR (n = 4 mice per group per time point). Limits of detection (LOD) indicated by the dotted lines. The dashed line indicates the limit of detection; viral RNA in the testis was not detected at any time point. N.D., not detected. Data are the mean ± s.e.m. Statistical significance was determined by two-way ANOVA with repeated measures followed by Sidak’s multiple-comparisons test (**a**), log-rank (Mantel–Cox) test (**b**), and one-way ANOVA with Dunnett’s multiple-comparisons test (**c**–**e**). The exact *P* values are indicated; **** *P* < 0.0001.

**Extended Data Fig. 3.**
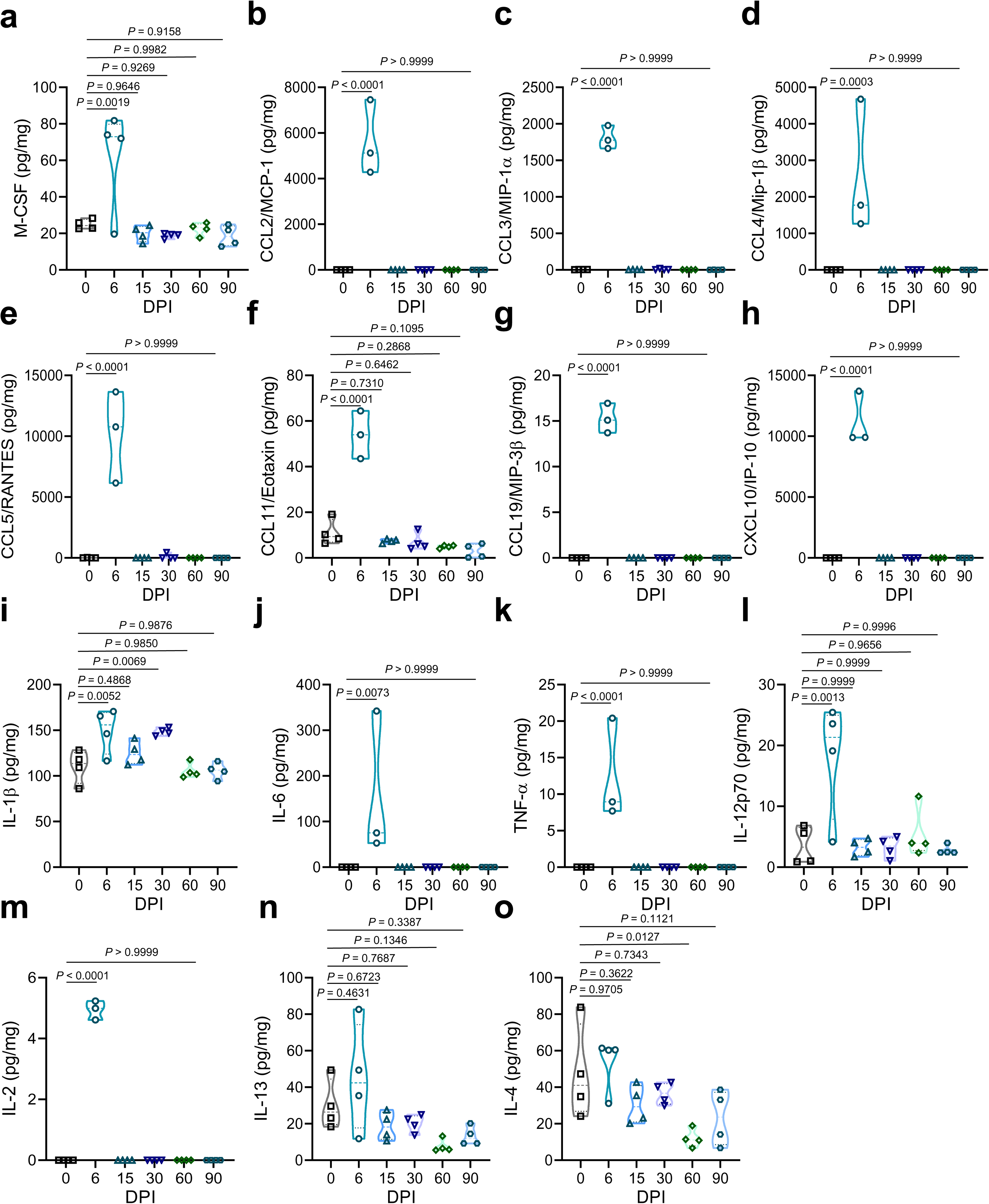
Multiplex profiling of cytokines and chemokines in whole-brain lysates from SARS-CoV-2-infected K18-hACE2 mice. **a–o**, Cytokine and chemokine protein concentrations measured in whole-brain lysates at 0 (mock), 6, 15, 30, 60 and 90 dpi using a multiplex bead-based array. Analytes include M-CSF (**a**), CCL2 (**b**), CCL3 (**c**), CCL4 (**d**), CCL5 (**e**), CCL11 (**f**), CCL19 (**g**), CXCL10 (**h**), IL-1β (**i**), IL-6 (**j**), TNF-α (**k**), IL-12p70 (**l**), IL-2 (**m**), IL-13 (**n**), and IL-4 (**o**). The data are presented as violin plots with overlaid individual data points (n = 4 per group at each time point; sporadic statistical outliers were excluded using the ROUT method, detailed in Methods). Non-detected samples are assigned a value of 0. Statistical significance was determined using one-way analysis of variance (ANOVA) with Dunnett’s multiple-comparisons test (each time point compared with 0 dpi). The exact *P* values are indicated.

**Extended Data Fig. 4.**
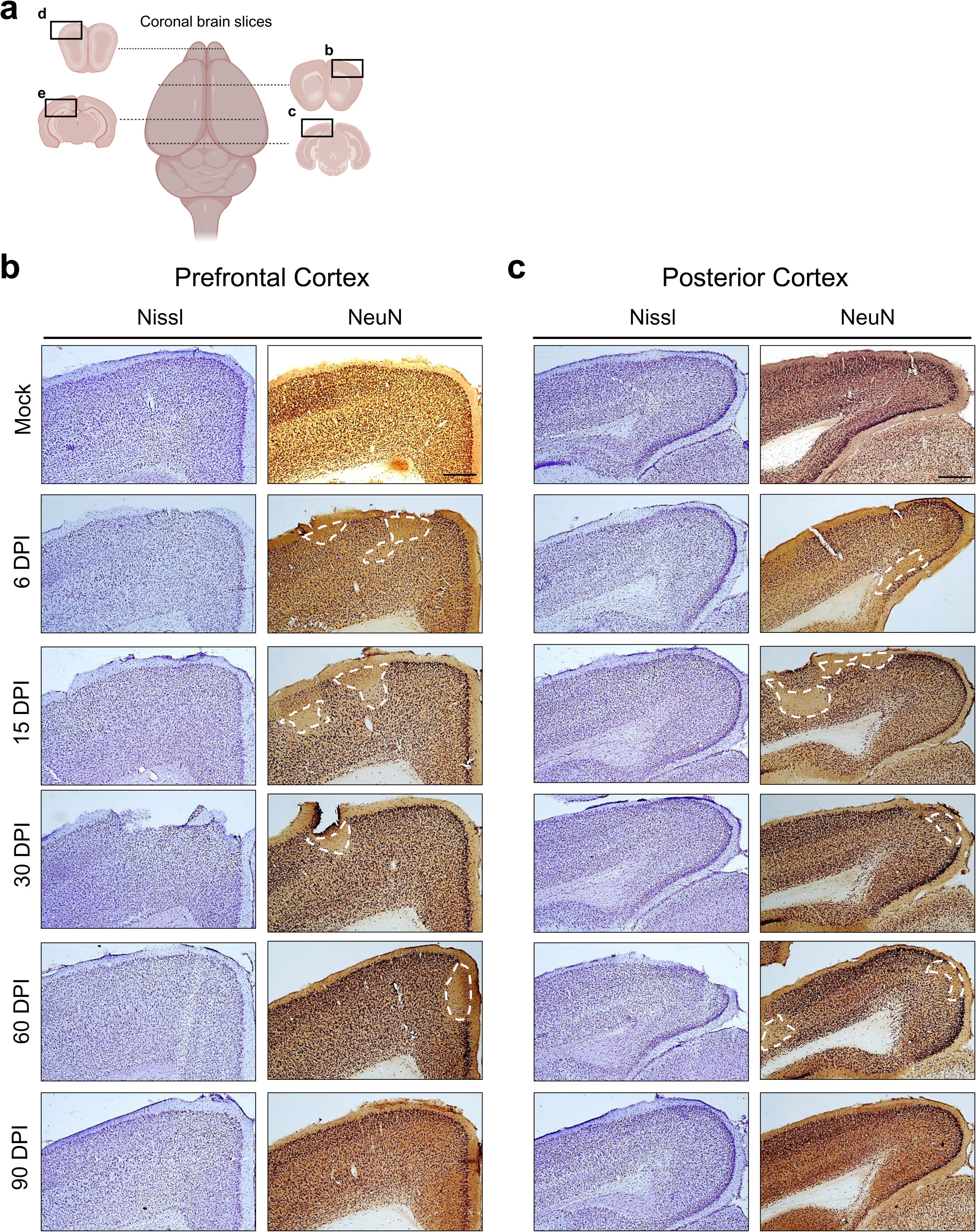
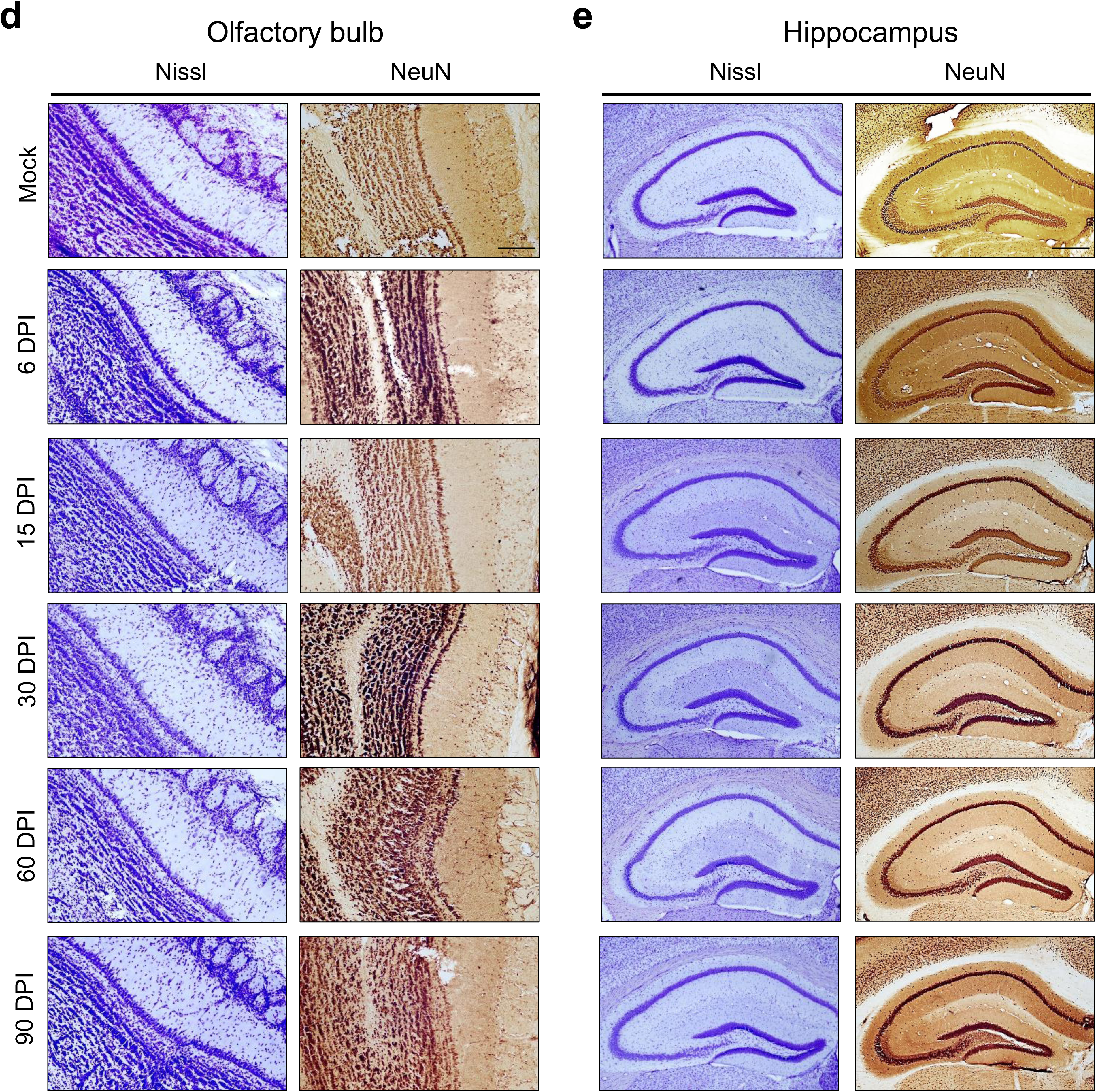
Persistent patchy reduction of neuronal NeuN expression in the cortex of K18-hACE2 mice following SARS-CoV-2 infection. **a**, Schematic of coronal brain sections indicating the anatomical locations of the prefrontal cortex (**b**), posterior cortex (**c**), olfactory bulb (**d**) and hippocampus (**e**). **b–e,** Representative images of Nissl (left) and NeuN (right) staining in the indicated brain regions from mock-infected and SARS-CoV-2-infected mice at 6, 15, 30, 60, and 90 dpi. White dashed lines demarcate focal areas of reduced NeuN immunoreactivity observed in the cortical regions (**b**,**c**), in contrast to preserved staining in the olfactory bulb (**d**) and hippocampus (**e**). Scale bars, 100 µm. The images are representative of n = 3-5 mice with similar results.

**Extended Data Fig. 5.**
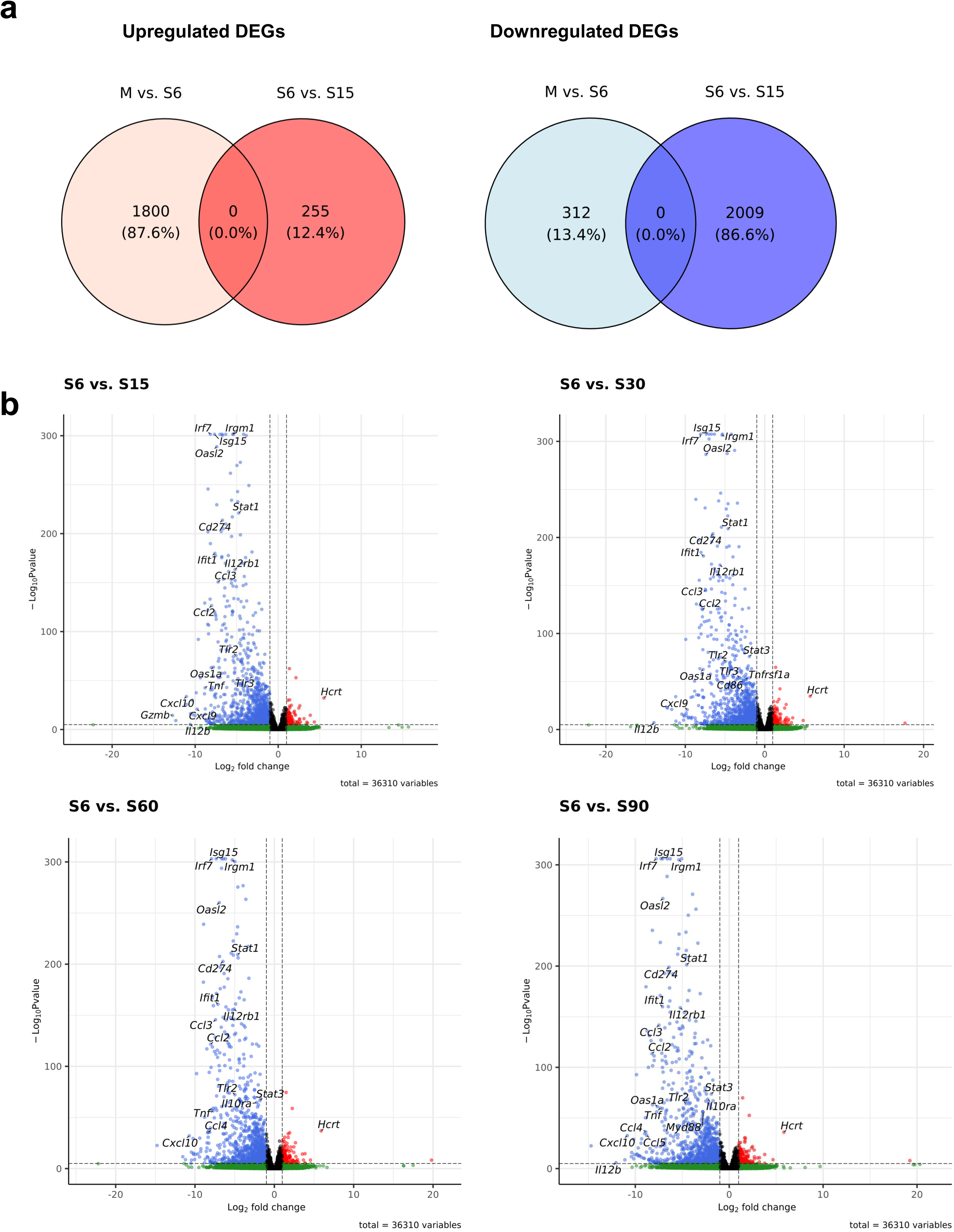
Transcriptomic profiling of SARS-CoV-2-infected K18-hACE2 mouse brains using RNA-seq. **a**, Venn diagrams illustrating the overlap of upregulated (left) and downregulated (right) differentially expressed genes (DEGs) between mock versus 6 days post-infection (dpi) (M vs. S6), and 6 dpi versus 15 dpi (S6 vs. S15) (M, mock; S6, 6 dpi; S15, 15 dpi). **b,** Volcano plots depicting differentially expressed genes (DEGs) for pairwise comparisons: 6 dpi versus 15 dpi, 6 dpi versus 30 dpi, 6 dpi versus 60 dpi, and 6 dpi versus 90 dpi. Key immune-related genes and Hcrt are labelled. Blue and red dots represent significantly downregulated and upregulated genes, respectively (log_2_ fold change > 1 or < −1; adjusted *P* < 0.05). The vertical dashed lines indicate the log_2_ fold-change cutoff, while the horizontal dashed line indicates the significance threshold.

**Extended Data Fig. 6.**
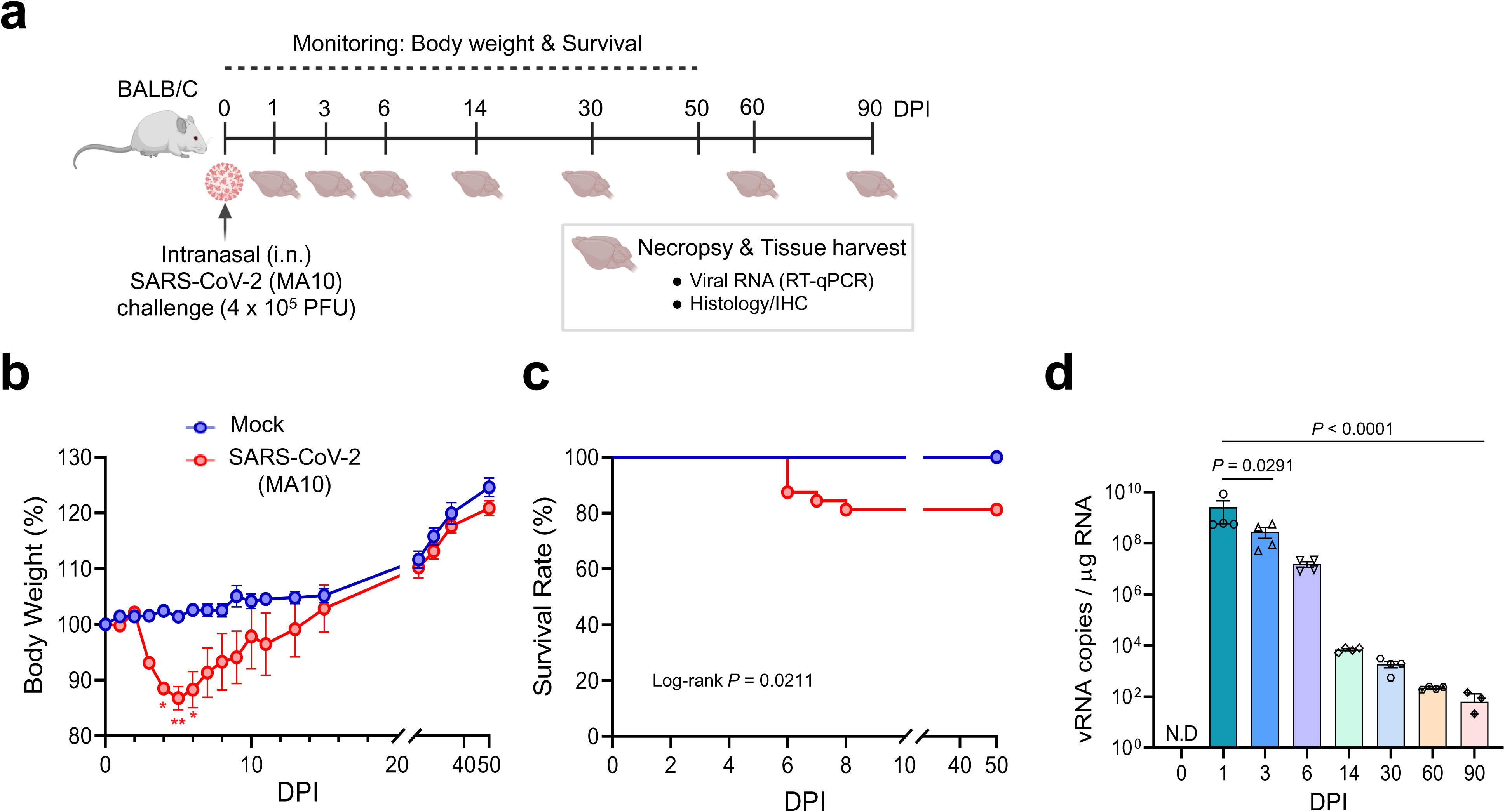
Clinical course and pulmonary viral RNA kinetics following MA10 SARS-CoV-2 infection in BALB/c mice. **a,** Experimental design. BALB/c mice were intranasally challenged with mouse-adapted SARS-CoV-2 (MA10; 4×10^5^ PFU) and monitored for up to 90 days post-infection (dpi); Tissues were collected for viral RNA quantification and histopathology at the indicated time points. **b,c,** Body weight change (**b**; mock n = 5; MA10 n = 12 mice, per group) and survival (**c**; mock n = 26, MA10 n = 32 mice per group) of MA10-infected mice compared with mock controls. **d,** Time-course quantification of lung viral RNA by RT–qPCR at the indicated time points (n = 4 mice per group per time point). The dashed line indicates the limit of detection. Data are the mean ± s.e.m. Statistical significance was determined using two-way ANOVA with repeated measures followed by Sidak’s multiple-comparison test (**b**), log-rank (Mantel–Cox) test (**c**) and one-way ANOVA with Dunnett’s multiple-comparisons test (**d**). The exact *P* values are indicated or denoted as \**P* < 0.05, \*\**P* < 0.01. N.D., not detected.

**Extended Data Fig. 7.**
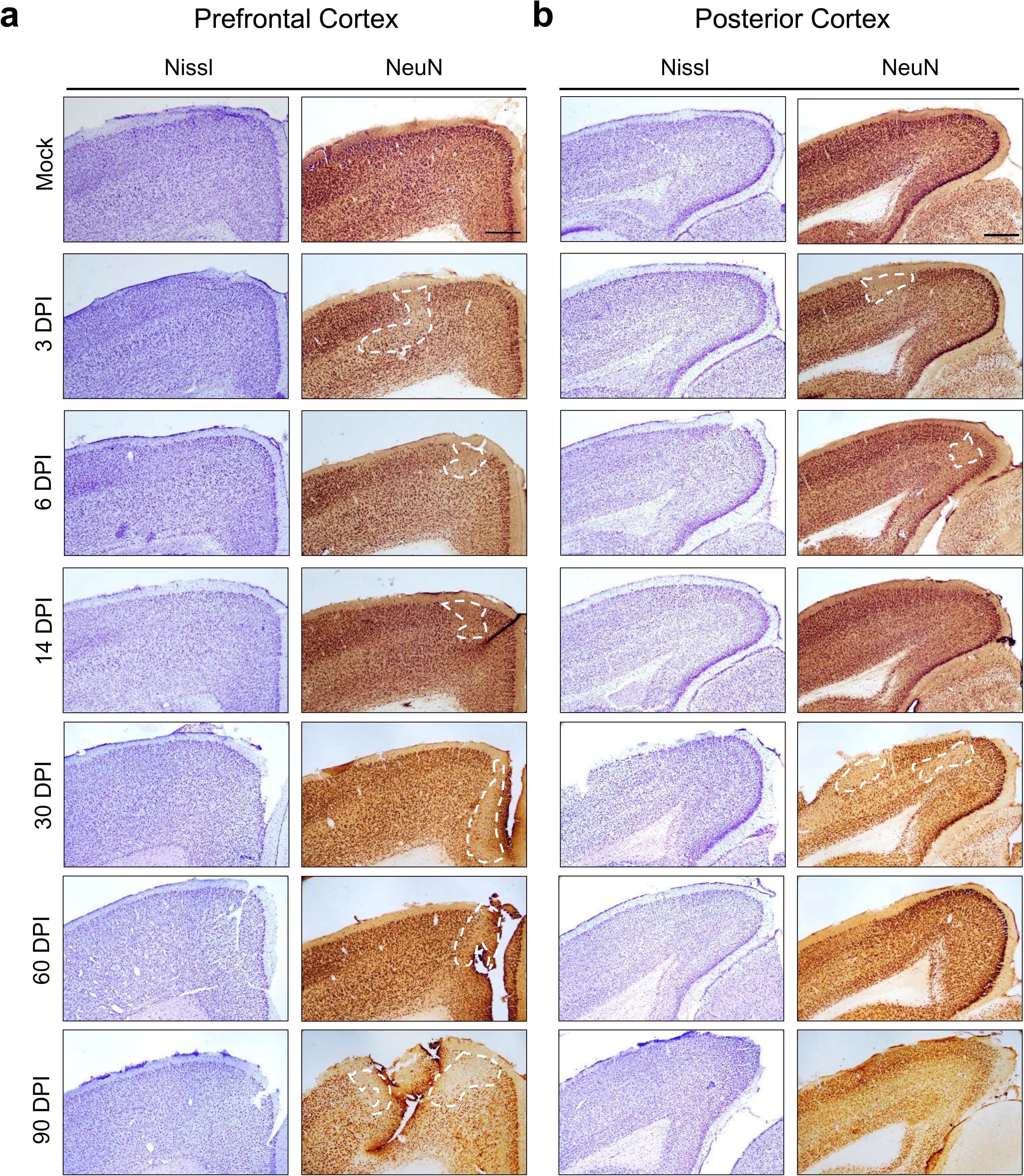
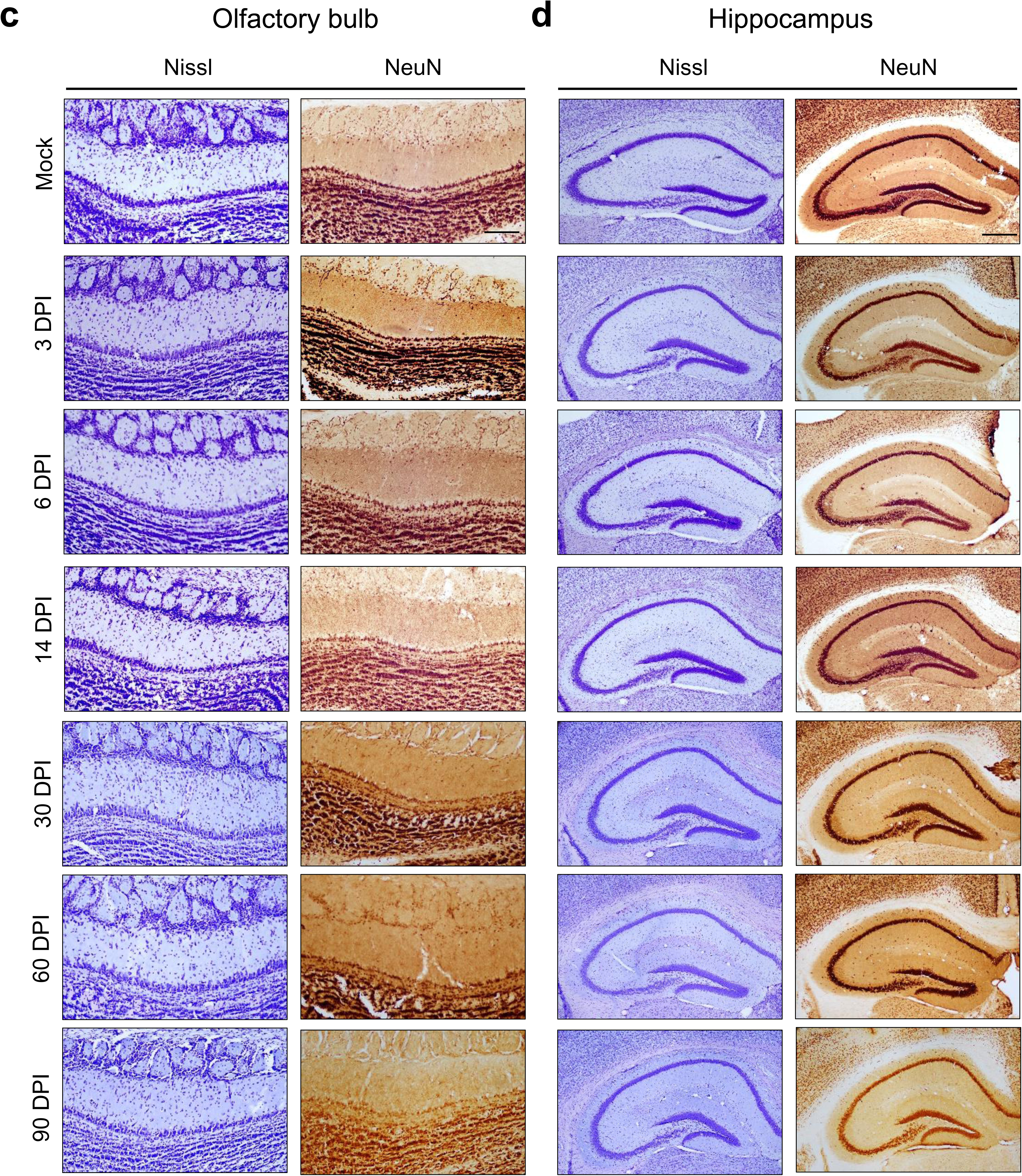
Persistent patchy reduction of cortical NeuN immunoreactivity in BALB/c mice following MA10 SARS-CoV-2 infection. **a,b**, Representative images of Nissl staining (left) and NeuN immunohistochemistry (right) of the prefrontal cortex (**a**) and posterior cortex (**b**) of mock- and MA10-infected BALB/c mice at 3, 6, 14, 30, 60, and 90 dpi. White dashed lines indicate the focal regions with reduced NeuN immunoreactivity. **c,d,** Representative images of Nissl staining and NeuN immunoreactivity in the olfactory bulb (**c**) and hippocampus (**d**) showing no overt reduction in NeuN immunoreactivity across the analysed time points. Scale bars: 100 µm (all panels). Images are representative of n = 3-4 independent biological replicates with similar results.

**Extended Data Fig. 8.**
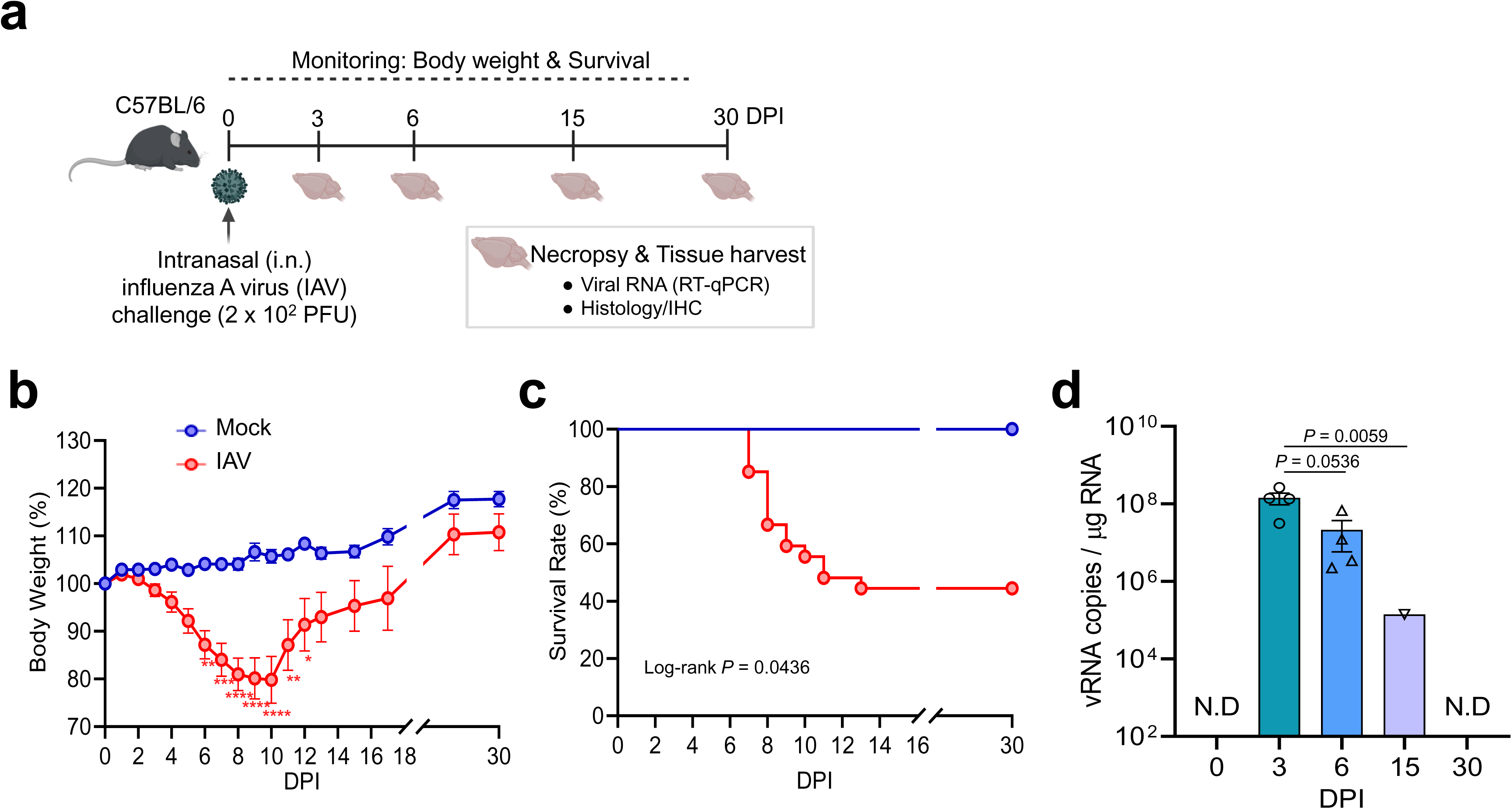
Long-term sequelae in influenza A (IAV)-infected C57BL/6 mice at 30 days post-infection. **a**, Schematic of the experimental design. C57BL/6 mice were intranasally challenged with the influenza A virus (IAV; PR8 strain, 2 × 10^2^ PFU). **b,c,** Body weight change (**b**; Mock, n = 5; IAV, n = 15) and survival analysis (**c**; Mock n = 5, IAV n = 27) of IAV-infected mice compared with mock controls. **d,** Time-course quantification of viral RNA loads in the lung at indicated time points using RT–qPCR (n = 4 mice per group). Data represent the mean ± s.e.m. Statistical significance was determined using two-way ANOVA with repeated measures followed by Sidak’s multiple comparison test (**b**), log-rank (Mantel–Cox) test (**c**) or one-way ANOVA with Tukey’s multiple comparison test (**d**). The exact *P* values are mentioned, or indicated by asterisks as follows: \**P* < 0.05, \*\**P* < 0.01, \*\*\**P* < 0.001, and \*\*\*\**P* < 0.0001. N.D., not detected.

**Extended Data Fig. 9.**
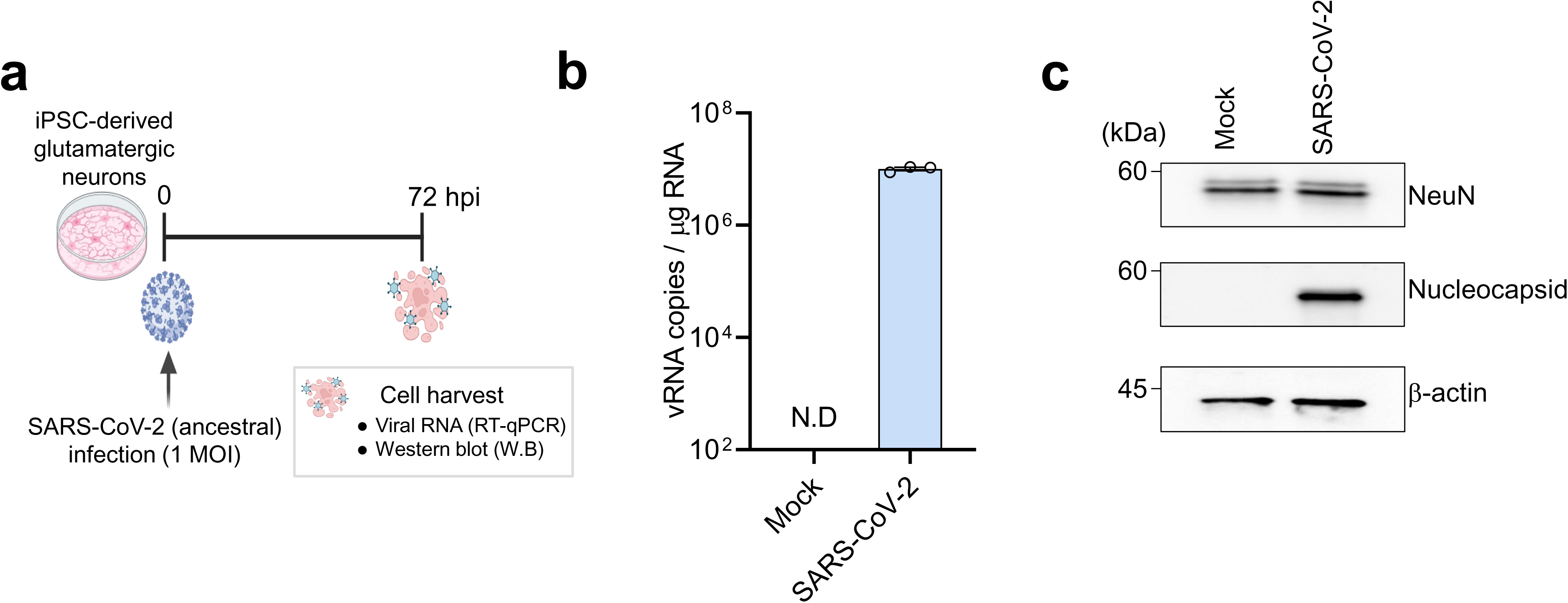
SARS-CoV-2 infection permits robust viral replication but does not deplete NeuN protein in human iPSC-derived glutamatergic neurones. **a**, Schematic of the experimental timeline. Human iPSC-derived glutamatergic neurones were infected with SARS-CoV-2 at a multiplicity of infection (MOI) of 1. Cells were harvested at 72 h post-infection (hpi) for analysis. **b,** Quantification of intracellular SARS-CoV-2 viral RNA loads using RT–qPCR at 72 hpi. Data are presented as viral RNA copies per µg of total RNA (n = 3 independent biological replicates). **c,** Western blot analysis of NeuN and SARS-CoV-2 Nucleocapsid (NP) protein expression in mock and SARS-CoV-2-infected neurones at 72 hpi. β- Actin was used as a loading control. Representative immunoblots are shown. Data represent the mean ± s.e.m.

**Table S1.**
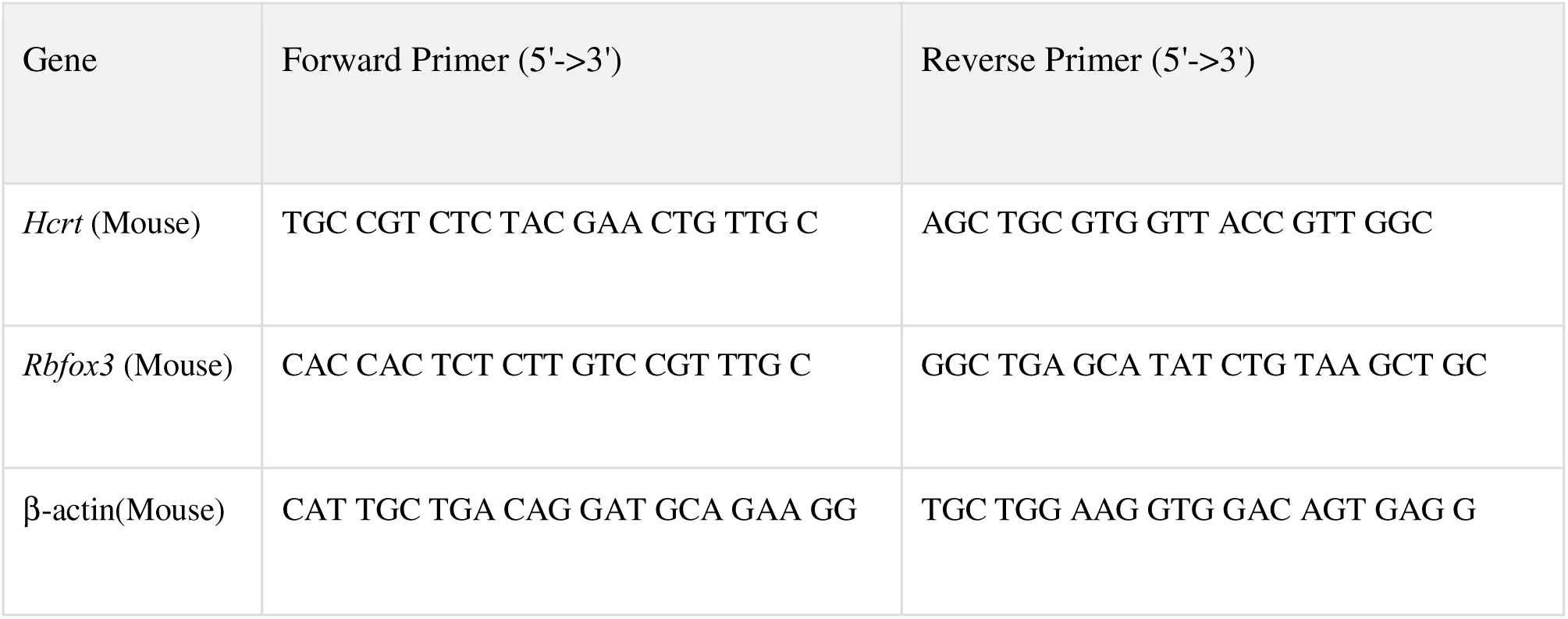
List of primers used for RT-qPCR.

